# A dicer-like3 protein affects paramutation at multiple loci in *Zea mays*

**DOI:** 10.1101/2023.09.29.560153

**Authors:** Ankur S. Narain, Irene T. Liao, Joy-El R.B. Talbot, Natalie C. Deans, Jay B. Hollick

## Abstract

Paramutation is a process by which meiotically-heritable gene regulation is altered by *trans*-homolog interactions. In *Zea mays*, genetic screens for functions maintaining paramutation-induced repressed states have identified loci encoding small RNA biogenesis components, thus implicating small RNAs in mediating these *trans*-homolog communications. Here we report that the *required to maintain repression5* locus encodes the sole dicer-like3 protein responsible for non-anther-specific 24-nucleotide RNA production. We found dicer-like3 is essential for mediating paramutation at the *booster1* locus and for the meiotic maintenance of transcriptionally repressed states at the *purple plant1* locus. Despite an expected role in mediating RNA-directed DNA methylation, we found 5-methylcytosine levels largely unchanged at multiple repetitive sequences in *dicer-like3* mutants, with minimal compensation from other small RNA sizes. The minor effects on plant heights and flowering time seen in the absence of dicer-like3 contrasts with other paramutation mutants and we highlight one specific allele repressed by RNA polymerase IV yet unaffected by dicer-like3 loss. These findings highlight diverse regulatory functions for individual components of 24-nucleotide biogenesis occurring in the grasses and support a working model in which this small RNA size class mediates *trans*-homolog interactions that drive meiotically-heritable changes in gene regulation.

## INTRODUCTION

Small RNAs (sRNAs) are widely recognized as key regulators of both eukaryotic gene expression (Borges and Martienssen 2015) and *trans*-generational epigenomic programming (Heard and Martienssen 2014), though their various modes of action in diverse species remain largely unknown. In fungi, plants, and animals, specific sized sRNAs in complex with argonaute-type effector proteins can direct DNA and/or chromatin modifications that dictate meiotically-heritable regulatory states (reviewed in Holoch and Moazed 2015; Matzke et al., 2015; Iwasaki et al., 2015). In *Zea mays* (maize), 24-nucleotide (nt) RNAs sourced from a plant-specific RNA polymerase IV (RNAP IV) (Erhard et al., 2009) are hypothesized to mediate paramutation (Alleman et al., 2006; Erhard et al., 2009; Chandler 2010; Arteagua-Vasquez et al., 2010; Belele et al., 2012; Hollick 2017), a process by which transcriptional repression of one allele (Patterson et al., 1993; Hollick et al., 2000; Sidorenko and Chandler 2008) is facilitated and stabilized in *trans* by the presence of locus-specific genomic features on the alternate homolog (Brink et al., 1960; Coe 1959; Hollick et al., 1995). This sRNA-centric hypothesis is largely based on current *Arabidopsis thaliana* (Arabidopsis) working models of RNA-directed DNA methylation (RdDM) (reviewed in Matzke and Mosher 2014; Matzke et al., 2015) in which RNAP IV-derived sRNAs (Li et al., 2015a; Blevins et al., 2015; Zhai et al., 2015a) of primarily, though not exclusively (Ye et al., 2016; Yang et al., 2016), 24-nt specify *de novo* cytosine methylation (5mC) in combination with nascent RNAP V transcripts. However, in both *Drosophila melanogaster* and *Caenorhabditis elegans*, model metazoans lacking 5mC modifications, certain germline-specific transgenes also exhibit paramutation-like behaviors that are dependent on cytoplasmic inheritance of sRNAs (deVanssay et al., 2012; Lutijn et al., 2012; Shirayama et al., 2012; Hermant et al., 2015; Sapetschnig et al., 2015).

Forward genetic approaches define at least fifteen *trans*-acting loci whose encoded functions affect paramutation-behaviors in maize (Dorweiler et al., 2000; Hollick and Chandler 2001; Hollick et al., 2005; Stonaker et al., 2009; Sidorenko et al., 2009; Sekhon et al., 2012; Sloan et al., 2014; Deans et al., 2020). Mutational screens using paramutant (transcriptionally repressed) *booster1* / *colored plant1* (*b1*) and *purple plant1* (*pl1*) alleles (*B1-Intense*; *B1-I* and *Pl1-Rhoades*; *Pl1-Rh*) identify *mediator of paramutation* (*mop*) and *required to maintain repression* (*rmr*) loci, respectively (Dorweiler et al., 2000; Hollick and Chandler 2001). Plant pigment serves as a quantitative readout of both *B1-I* (Patterson et al., 1993) and *Pl1-Rh* (Hollick et al., 2000) expression because B1 and PL1 are transcriptional activators of anthocyanin biosynthetic genes (Dooner et al., 1991). *B1-I* and *Pl1-Rh* condition strong pigmentation in *B-I* and *Pl-Rh* reference states, but derivative paramutant *B′* and *Pl′* states confer relatively weak and variegated anthocyanin deposition (Emerson 1921; Coe 1959; Hollick et al., 1995). In homozygous conditions, recessive *mop* and *rmr* mutations intensify pigments of *B′* / *B′* or *Pl′* / *Pl′* plants to *B-I-* or *Pl-Rh*-like levels respectively (Dorweiler et al., 2000; Hollick and Chandler 2001).

All but two of the identified MOP and RMR proteins are known constituents of macromolecular complexes (Haag et al., 2014) including 1) both catalytic subunits of RNAP IVa, one of at least two distinct grass-specific RNAP IV subtypes (Erhard et al., 2009; Stonaker et al., 2009; Sidorenko et al., 2009; Sloan et al., 2014; Huang et al., 2015), 2) MOP1, an RNA-dependent RNA polymerase sharing similarity with Arabidopsis RNA DEPENDENT RNA POLYMERASE 2 (RDR2) (Alleman et al., 2006; Woodhouse et al., 2006), and 3) RMR1 (Hale et al., 2007), a Rad54-like ATPase defining a CLASSY (CLSY)-related clade of Arabidopsis proteins (Smith et al., 2007). Outside these complexes, RMR2 represents the founding member of a large class of plant-specific proteins with no recognizable structured domains (Barbour et al., 2012), and RMR12 is a chromodomain helicase DNA-binding 3 (CHD3) protein orthologous to the Arabidopsis PICKLE nucleosome remodeler (Deans et al., 2020). Because all these maize proteins are required for normal 24-nt RNA biogenesis (Alleman et al., 2006; Hale et al., 2007; Nobuta et al., 2008; Erhard et al., 2009; Stonaker et al., 2009; Sidorenko et al., 2009; Barbour et al., 2012) or defining genomic locations thereof (Deans et al., 2020), it is proposed that paramutant states are maintained by an RdDM-type mechanism.

The potential of 24-nt RNAs to mediate *trans*-homolog changes in chromatin structures also provides a working model to explain one of the hallmarks of paramutation behaviors (Greaves et al., 2012; Gouli et al., 2016). *B-I* and *Pl-Rh* invariably change to *B′* and *Pl′* in *B′* / *B-I*, or *Pl′* / *Pl-Rh* heterozygotes respectively (Coe 1959; Hollick et al., 1995). These “induced” paramutations require both MOP1 (Dorweiler et al., 2000; Hollick et al., 2005) and the largest catalytic subunit, rna polymerase d1 (RPD1/RMR6/MOP3; orthologous with Arabidopsis NUCLEAR RNA POLYMERASE D1, NRPD1), of both RNAP IV subtypes. Paramutation occurring in *B-I / B′* plants can also be inhibited in *rmr7-1* mutants having an early stop codon in the gene encoding RP(D/E)2a (orthologous to Arabidopsis NUCLEAR RNA POLYMERASE D2, NRPD2) (Stonaker et al., 2009), one of at least two RNAP IV and three RNAP V second-largest subunits (Haag et al., 2014). These findings support a model in which *B′* and *Pl′* states provide allele-specific RNAP IV-derived sRNAs that then direct conversion of *B-I* and *Pl-Rh* to *B′* and *Pl′* respectively.

Sequences responsible for inducing paramutation at both *B1-I* and *Pl1-Rh* appear to be coincident with distal transcriptional enhancers (Stam et al., 2002; Erhard et al., 2013). The 3′ *Pl1-Rh* sequences remain undefined, but the *B1-I* enhancer is contained within a tandem array of seven near-identical 853bp repeats (hepta-repeat) found approximately 100 Kb 5′ of the *b1* coding region (Stam et al., 2002; Belele et al., 2013) that loop to the *b1* promoter in frequencies correlated with *b1* transcription rates (Patterson et al., 1993; Louwers et al., 2009). In contrast to the expectations of a simple RdDM-type model, both *B-I* and *B′* – as well as a recessive *b1* allele that has no paramutation-type behaviors – produce apparently equivalent MOP1- dependent 24-nt RNAs representing the distal hepta-repeat (Arteagua-Vasquez et al., 2010). It is also unclear how *Pl′* / *Pl-Rh* changes to *Pl′* / *Pl′* in the absence of RMR1-dependent 24-nt RNAs (Hale et al., 2007). Additionally, even though *Pl′* states can revert back to meiotically-heritable *Pl-Rh* in some homozygous *mop* and *rmr* mutants (Dorweiler et al., 2000; Hollick and Chandler 2001; Hollick et al., 2005; Hale et al., 2007), *B′* states do not (Dorweiler et al., 2000; Hollick et al., 2005; Stonaker et al., 2009; Sidorenko et al., 2009). Thus, the current RdDM-type working model for maize paramutation requires further testing and potential refinement.

The canonical Arabidopsis RdDM model maintains a feed-forward loop of sRNA biogenesis, 5mC, histone H3 lysine 9 (H3K9) methylation, and RNAP IV recruitment (reviewed in Matzke and Mosher 2014; Matzke et al., 2015). RNAP IV / RDR2 / CLSY1/2 complexes are recruited to chromatin via an adapter protein (DNA-BINDING TRANSCRIPTION FACTOR 1/SAWADEE HOMEODOMAIN HOMOLOG1) that binds H3 N-terminal tails having unmethylated K4 but methylated K9 modifications (Law et al., 2013; Zhang et al., 2013; Zhao et al., 2022). Recently, it has been found that RNAP IV / CLSY3/4 complexes in developing ovules are recruited by an alternate mechanism associated with an AT-rich sequence motif (Zhao et al., 2022). RNAP IV produces short 25-50-nt primary transcripts (Blevins et al., 2015; Zhai et al., 2015a) that are made double-stranded by RDR2 (Li et al., 2015a) and primarily processed by the DICER-LIKE 3 (DCL3) endonuclease into 24-nt/24-nt or 24-nt/23-nt duplexes (Loffer et al., 2022) from which a single-stranded guide RNA is loaded/stabilized in ARGONAUTE 4 (AGO4). Arabidopsis has four DCL proteins dedicated to generating specific sized cleavage products (Xie et al., 2004; Kasschau et al., 2007) having some semi-redundant functions (Henderson et al., 2006; Fahlgren et al., 2006; Schmitz et al., 2007), but *in vitro*, DCL3 prefers small double-stranded (ds) RNA precursors (Nagano et al. 2014) like those generated from RNAP IV / RDR2 (Li et al., 2015a; Blevins et al., 2015; Zhai et al., 2015a). A DCL-independent pathway involving exonuclease trimming can also generate 24-nt RNA / AGO4 complexes that maintain RdDM at a subset of genomic targets (Ye et al. 2016; Yang et al. 2016). Recruitment of a *de novo* DNMT3-like DNA methyltransferase is targeted by AGO4 interactions with the C-terminal domain of the RNAP V largest subunit and by presumed base pairing between the guide RNA and nascent RNAs (Zhong et al., 2014; Sigman et al., 2021), and/or the DNA template (Lahmy et al., 2016). RNAP V is guided to regions enriched in preexisting 5mC via two 5mC-binding proteins, SUPPRESSOR OF VARIEGATION (3-9) HOMOLOG (SUVH) 2 and SUVH9 (Johnson et al., 2014; Liu et al., 2014), or through AGO4-based recruitment to nascent RNAP II RNAs (Sigman et al., 2021). The CHROMOMETHYLASE 3 (CMT3), in combination with histone methyltransferase SUVH4 / KRYPTONITE (KYP), provide a self-reinforcing maintenance of 5mC and H3K9me2 marks (Du et al., 2014). Even with many of these plant-specific molecules, the basic feed-forward loop of sRNA-directed chromatin modifications affecting proper RNA transcription and mRNA processing appears to be a highly conserved eukaryotic mechanism (reviewed in Holoch and Moazed 2015).

Given the diversification of core RdDM-related proteins found within the grasses (Qian et al., 2011; Huang et al., 2015; Zhang et al., 2015), including the aforementioned RNAP IV and V subtypes, their uses in maize may be different from those of a canonical Arabidopsis pathway. Maize also lacks a CMT2 ortholog (Jiao et al., 2017; Zemach et al., 2013) which, in Arabidopsis, maintains 5mC patterns at ∼70% of all cytosines found in CHH (H = A, T, or C) contexts (Stroud et al., 2013; Zemach et al., 2013). Maize and *Oryza sativa* (rice) also have five dicer-like genes (Qian et al., 2011). In maize, DCL1 processes microRNAs (Thompson et al., 2014) and some *trans*-acting siRNAs redundantly with DCL4 (Petch et al., 2015), and the grass-specific DCL5 (Margis et al., 2006; Qian et al., 2011; Song et al., 2012) produces phased 24-nt RNAs (phasiRNAs) representing 176 genome-dispersed loci in the tapetal cells of developing anthers (Zhai et al., 2015b). The functions of maize DCL2 and DCL3 remain unknown.

Here we show that four recessive ethyl methanesulfonate (ems)-induced mutations define the novel *rmr5* locus with functions required for maintaining the meiotically-heritable *Pl′* state and for mediating paramutation in *B′* / *B-I* heterozygotes. Positional mapping and DNA sequencing led to the identification of *rmr5* as a single-copy gene encoding the sole ZmDCL3 enzyme. Mutant sRNA profiles are consistent with the ZmDCL3 assignment, but comparisons show many 5mC patterns are maintained independent of ZmDCL3, even though few sRNA size classes appear elevated in *rmr5* mutants. Minor delays in flowering time and slightly reduced stature were documented in *rmr5* mutants, but unlike rice *dcl3* RNAi-knock down lines (Wei et al., 2013), no major developmental defects were observed. Comparisons with *rpd1*/*rmr6* mutant profiles identified ZmDCL3-independent effects that implicate additional RNAP IV roles outside of a canonical RdDM mechanism. The finding that ZmDCL3 affects paramutation at multiple loci supports working models in which 24-nt RNAs mediate *trans*-homolog interactions.

## RESULTS

### The *rmr5* Locus is Defined by Recessive *rmr*-type Mutations

Through an ongoing genetic screen for ems-induced *rmr-*type mutations conferring darkly-colored anthers to *Pl′* / *Pl′* plants (Hollick and Chandler 2001; Hale et al., 2007), we identified four single-locus recessive (Supplemental Table 1 and Supplemental Methods) mutations that failed to genetically complement each other, but complemented mutations defining previously characterized *rpd1* (*rmr6*), *rp(d/e)2a* (*rmr7*), *rmr1*, *rmr2,* and *mop1* loci (Supplemental Table 2 and Supplemental Methods). Additionally, none of the mutants had developmental defects diagnostic of *chd3a* (*rmr12*) mutations (Deans et al., 2020). These results indicate that a novel locus, *required to maintain repression5* (*rmr5*), is defined by the mutations found within *rmr5-1*, *rmr5-2*, *rmr5-3*, and *rmr5-4* alleles.

### *Rmr5* Maintains Paramutagenic *Pl′* **States**

Homozygous mutants derived from self-pollinating *Rmr5* / *rmr5* ; *Pl′* / *Pl′* heterozygotes have darkly-pigmented anthers, indistinguishable from *Pl-Rh* / *Pl-Rh* genotypes (Supplemental Table 1). To evaluate the epigenetic state(s) of *Pl1-Rh* alleles transmitted from these mutants, pollen from one individual M_2_ *rmr5-3* homozygote was divided and applied to related *Pl′* / *Pl′* and *Pl-Rh* / *Pl-Rh* plants representing three distinct inbred lines (A619, W23, A632) and the anther pigmentation of the resulting progenies were visually scored on a 1 – 7 graded scale (Hollick et al., 1995). All progeny of *Pl′* / *Pl′* females had low anther color scores (1 or 2) (Table 1) implying that the *rmr5-3* mutation is completely recessive and that *Pl1-Rh* alleles transmitted from this individual *rmr5-3* mutant are not recalcitrant to subsequent paramutation. In contrast, anther color scores of progeny derived from crosses to 2 of 3 different *Pl-Rh* / *Pl-Rh* plants (A619 and W23) ranged from 1 to 7 (Table 1) implying that some sperm cells transmitted *Pl1-Rh* alleles that had reverted from *Pl′* to *Pl-Rh*-like states in this single homozygous *rmr5-3* M_2_ plant. Progeny of the A632 *Pl-Rh* / *Pl-Rh* cross, however, all had low anther color scores. These inbred line-specific differences in detecting apparent reversion events mirror observations that *Pl-Rh* states are most stable in both A619 and W23 lines yet relatively unstable in A632 (Hollick et al., 2005). However, six of twenty-six (26%) individuals from the 070181 progeny derived by crossing a A632 *Pl-Rh* / *Pl-Rh* female by an *rmr5-4* M_2_ mutant had anther color scores of 7 illustrating that reversion events could be detected using all three *Pl-Rh* / *Pl-Rh* lines.

**Table 1.** Paramutagenicity of *Pl1-Rh* Following Transmission from *rmr5-3* Mutants.

| Testcross progeny of *Rmr5* / *Rmr5* ; *PI1-Rh* / *PI1-Rh* X *rmr5-3* / *rmr5-3* ; *PI'* / *PI'* crosses |  |  |  |  |  |  |  |  |  |
| --- | --- | --- | --- | --- | --- | --- | --- | --- | --- |
| Progeny Identifier | Female genotype | Mutant generation | No. of progeny individuals with specific ACS |  |  |  |  |  |  |
|  |  |  | 1 | 2 | 3 | 4 | 5 | 6 | 7 |
| 060630 | A619 *PI'* | M2 | 3 | 3 | 0 | 0 | 0 | 0 | 0 |
| 060617 | A619 *PI-Rh* | M2 | 1 | 7 | 1 | 0 | 0 | 0 | 1 |
| 060392 | W23 *PI'* | M2 | 7 | 5 | 0 | 0 | 0 | 0 | 0 |
| 060431 | W23 *PI-Rh* | M2 | 1 | 6 | 0 | 5 | 1 | 1 | 0 |
| 060420 | A632 *PI'* | M2 | 7 | 1 | 0 | 0 | 0 | 0 | 0 |
| 060419 | A632 *PI-Rh* | M2 | 7 | 8 | 0 | 0 | 0 | 0 | 0 |
| 063295 | A619 *PI-Rh* | M2S1* | 0 | 1 | 0 | 1 | 0 | 2 | 5 |
| 064108 | W23 *PI-Rh* | M2S1* | 2 | 1 | 1 | 3 | 4 | 2 | 12 |
| 151053 | A619 *PI-Rh* | M2S4* | 0 | 0 | 0 | 0 | 0 | 0 | 15 |
| 151060 | A619 *PI-Rh* | M2S4* | 0 | 0 | 0 | 0 | 0 | 0 | 11 |

\* M2S1 and M2S4 derive from the same M2 individual used in original reversion tests.

To evaluate whether loss of *rmr5* function over multiple generations led to decreased stability of the *Pl′* state, pollen from a single M_2_S_1_ *rmr5-3* mutant was also divided and applied to both A619 and W23 *Pl-Rh* / *Pl-Rh* representatives. Approximately half of all progeny from both crosses had anther color scores of 7 indistinguishable from *Pl-Rh* / *Pl-Rh* genotypes and few (2/9 and 7/25) had scores of 1 to 4 representing *Pl′* / *Pl′* types (Table 1). Later crosses between A619 *Pl-Rh* / *Pl-Rh* and two independent M_2_S_4_ *rmr5-3* mutants gave rise to progeny of exclusively ACS 7 type (26/26) (Table 1). These observations indicate that *Rmr5* functions act somatically to maintain repressed *Pl′* states and that meiotically-heritable reversions of *Pl′* to non-paramutagenic *Pl-Rh* states occur with persistent *trans*-generational absence of *Rmr5* function. These behaviors are similar to those detailed previously for specific *mop1* (Dorweiler et al., 2000), *rmr1* (Barbour et al., 2007), and *rp(d/e)2a* / *rmr7* (Stonaker et al., 2009) mutations.

### *Rmr5* is Required for *b1* Paramutation

In contrast to *Pl′, B′* states are irreversible when transmitted from *B′* / *B-I* heterozygotes (Coe, 1966; Patterson et al., 1993; Chandler, 2000). We could therefore reliably evaluate whether *Rmr5* function was essential for the induction of *b1* paramutation by synthesizing *rmr5-1* / *rmr5-1* ; *B′* / *B-I* plants and assessing the plant colors of testcross progeny (Methods). Near isogenic *rmr5-1* / *Rmr5-W23* ; *b1* / *B′* and *rmr5-1* / *Rmr5-W23* ; *b1* / *B-I* F_1_ plants were crossed to generate the desired genotype. By testcrossing F_2_ plants having dark anthers (*rmr5-1* / *rmr5-1*) and any stem/sheath color (*b1* / *B′*, *b1* / *B-I*, or *B′* / *B-I*) by a recessive *b1* / *b1* pollen source, we could retrospectively define the female *b1* genotype by subsequent examination of progeny plant colors (Table 2). The single *b1* / *B′* ; *rmr5-1* / *rmr5-1* F_2_ individual had a plant color indistinguishable from *B′* indicating that *Rmr5* is not required for the somatic repression diagnostic of *B′* states. In contrast, all three *B′* / *B-I* ; *rmr5-1* / *rmr5-1* plants had a clear *B-I*-like phenotype indicating that *trans*-homolog silencing was impaired. Among the three testcross progenies derived from *B′* / *B-I* ; *rmr5-1* / *rmr5-1* plants, each one had nearly equal numbers of *B′*- and *B-I*-type individuals (Table 2) consistent with the interpretation that normal *Rmr5* function is required to mediate or establish paramutation at *b1*. This role of *Rmr5* in the paramutation mechanism operating at multiple maize loci motivated efforts to identify the molecular nature of the *rmr5* locus.

**Table 2.** Testcross Evaluating *Rmr5* Effects on *b1* Paramutation.

| Testcross progeny of *b1* / *b1* X (*b1* or *B1-I*) / (*b1* or *B'*) ; *rmr5-1* / *rmr5-1* crosses |  |  |  |  |  |
| --- | --- | --- | --- | --- | --- |
| Mutant parent ID | Mutant parent phenotype | Progeny |  |  |  |
|  |  | Identifier | Plant phenotype |  |  |
|  |  |  | *B-I*-type | *B'*-type | colorless |
| 10-563-1 | B-I | 101856 | 5 | 0 | 4 |
| 10-563-3 | B-I | 101854 | 3 | 0 | 3 |
| **10-563-18** | **B-I** | **101853** | **21** | **22** | **0** |
| **10-563-21** | **B-I** | **101857** | **3** | **6** | **0** |
| 10-563-23 | B' | 101852 | 0 | 6 | 6 |
| 10-563-33 | B-I | 101862 | 0 | 0 | 1 |
| **10-563-41** | **B-I** | **101855** | **26** | **14** | **0** |
| 10-563-50 | B-I | 101860 | 3 | 0 | 2 |
| 10-563-62 | B-I | 101861 | 0 | 0 | 3 |
| 10-563-67 | B-I | 101859 | 3 | 0 | 3 |
| 10-563-69 | B-I | 101858 | 3 | 0 | 5 |

Bolded entries represent analyses of *rmr5-1* mutants having a *B-I* / *B'* genotype.

### The *rmr5* Locus Maps to Chromosome *3L*

Because *rmr5* mutations were induced in A619 pollen crossed to a polymorphic A632 recipient (Hale et al., 2007), we used Simple Sequence Length Polymorphisms (SSLP) to locate the *rmr5* locus to a 6 Mb region of chromosome *3L*. Genomic DNA was isolated from 177 *rmr5* mutant BC_1_F_2_ seedlings (56 *rmr5-1*, 81 *rmr5-2*, and 40 *rmr5-3*) initially identified by their fully-colored leaf sheath phenotypes and subsequently confirmed at flowering as having darkly-colored anthers. A total of 229 oligonucleotide primer sets identifying A619/A632 SSLPs throughout the maize genome were selected or developed for these mapping efforts (Supplemental Table 3). Bulked segregant analysis identified three *3L* SSLP markers (bnlg1505, umc2265, and umc1593) biased toward A619 character (Supplemental Figure 1). Additional, local SSLP markers were developed and used to genotype genomic DNA samples from 98 individual *rmr5* mutants that narrowed the *rmr5* interval between 159 and 165 Mb relative to the ZmB73 RefGen_v3 containing 130 genes (Supplemental Table 4).

### Mutations Support a *dcl3* Gene Model Representing the *rmr5* Locus

Within the mapped interval, one gene model (GRMZM5G814985, ZmB73 RefGen_v3; Zm00001d042432, ZmB73 AGPv4) is predicted to encode a protein containing all features typical of a dicer enzyme (MacRae et al., 2007) including two presumed catalytic domains typical of all RNase III proteins, an N-terminal ATPase/DEAD-type helicase, a dsRNA binding motif, and a PAZ domain (Figure 1A). This protein is the putative *Zea mays* ortholog (previously annotated as DCL104) of Arabidopsis DICER-LIKE3 encoded by AT3G43920 (Figure 1B). This *dcl104* gene model was considered a strong *rmr5* candidate because Arabidopsis DCL3 is the RNase III-type endonuclease responsible for generating 24-nt RNAs from RNAP IV / RDR2 dsRNAs (Xie et al., 2004; Kasschau et al., 2007) and both MOP1 / ZmRDR2 and RPD1 are required for facilitating *b1* paramutation (Dorweiler et al., 2000; Hollick et al., 2005). Sequence obtained from PCR amplicons (Supplemental Table 5) representing the predicted 24 GRMZM5G814985 exons from *rmr5-2* / *rmr5-2* genomic DNA identified a single C to T transition-type nonsense mutation in exon 4, supporting this candidate gene assignment. Single transition mutations, typical of ems-mutagenesis, creating other nonsense codons within the same gene model were also found in the other three *rmr5* mutant alleles (Figure 1A) strongly indicating that the *rmr5* locus encodes DCL104. B73 inbred reference sequence obtained from cDNAs (Supplemental Table 6) representing total RNA from a 4.5cm immature cob, validated the exon and intron structure of this 9511bp (ATG to TGA) gene model encoding a 1659 amino acid polypeptide having conserved domain organizations typifying other DICER-LIKE 3 proteins (Margis et al., 2006; Mukherjee et al., 2013)(see Methods and Supplemental Table 7).

**Figure 1.**
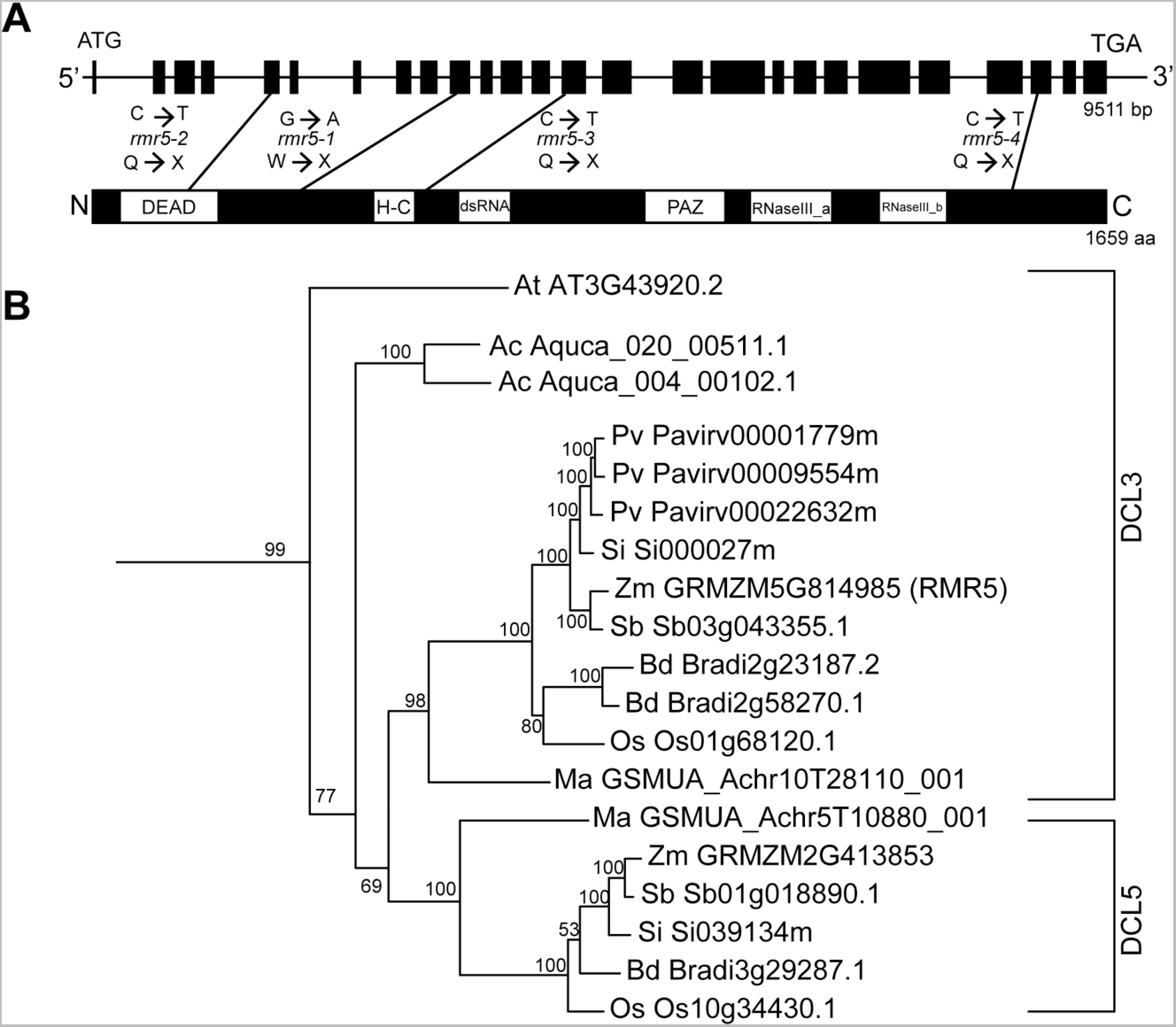
*rmr5* Candidate Gene Model and Phylogeny. **(A)** Schematic of the *dcl104* gene model (above) illustrating exons (black rectangles) with lines connecting the location of specific ems-induced mutations and the resulting nonsense codons in the predicted protein model (below). Protein model highlights six domains diagramed after the annotations from the Pfam search algorithm: DEAD (DEAD/DEAH box helicase), H-C (Helicase-conserved C-terminal domain), dsRNA (double stranded RNA binding domain), PAZ (PAZ domain), and RNaseIII. **(B)** Maximum likelihood relationships based on alignments for representative grasses: maize (Zm), sorghum (*Sorghum bicolor;* Sb), rice (*Oryza sativa;* Os), switchgrass (*Panicum virgatum;* Pv), foxtail millet (*Setaria italic;* Si), *Musa acuminata* (Ma), and *Brachypodium distachyon* (Bd); the eudicot *Aquilegia coerulea* (Ac) and *Arabidopsis thaliana* (At) outgroup. Branch lengths correspond to the indicated bootstrap values (n=100). The DCL3 and DCL5 clades are indicated.

A maximum likelihood analysis among representative monocots and the eudicots *Aquilegia coerulea* and Arabidopsis highlights the relative diversity of DCL3-type proteins found in grasses (Figure 1B)(Margis et al., 2006; Mukherjee et al 2013). The B73 genome has one other gene model encoding a similar dicer-like3 protein (DCL102; GRMZM2G413853, RefGen_v3; Zm00001d032655, AGPv4) clustering with rice DCL3b (Figure 1B), an inflorescence-restricted protein generating 24-nt phasiRNAs (Song et al., 2012) similar to those produced during maize microsporogenesis (Zhai et al., 2015b). Based on the amino acid divergence between rice OsDCL3a and OsDCL3b (∼57% similar), the highly divergent dsRNA binding motif of DCL3b representatives (Margis et al., 2006), retention of both DCL3a and 3b types in representative monocots (Figure 1B), and neofunctionalized phasiRNA biogenesis, we concur with the suggestion of Margis et al. (2006) and usage by Zhai et al. (2015b) that previously described DCL3b proteins be identified as DCL5. All monocots, with the apparent exception of switchgrass (*Panicum virgatum*), have a single gene encoding DCL5 (Supplemental Table 8) placing the presumed specialization of DCL5 function prior to the *Zingiberales* and *Poales* divergence. Despite the relatively recent maize tetraploidy (Gaut and Doebley 1997), only single examples of genes encoding DCL1, 2, 3, 4, and 5 remain (Supplemental Table 8). This finding implies that the duplicate ZmDCL genes have been lost through fractionation, possibly involving short intrachromosomal deletions (Woodhouse et al., 2010). In contrast, *Brachypodium distachyon*, *Musa acuminate*, and *Panicum virgatum* have all retained one or more DCL3-encoding duplicate (Figure 1B).

### RMR5 is Required for 24-nt RNA Accumulation

Consistent with *rmr5* representing the sole locus encoding a dicer-like3 enzyme, we found that a sRNA library produced from a 4.5cm immature cob of an *rmr5-2* homozygote was depleted for 24-nt reads (24mers) relative to that of a heterozygous sibling (6% vs 73% of total reads) (Figure 2; Supplemental Tables 9 and 10). Normalizing these datasets to the levels of either 21- or 22-nt species indicate that greater than 95% of all 24mers, whether they represent all, or exclusively unique, genome sequences (Figures 2A, 2B) are *Rmr5* dependent. These data confirm the assignment of the *Zea mays* DCL3 ortholog (ZmDCL3) as being encoded by *rmr5/dcl104*, hereafter synonymously named *dcl3*. Mutant alleles *rmr5-1*, *2*, *3*, and *4* are subsequently referred to as *dcl3-1, 2, 3*, and *4* respectively.

**Figure 2.**
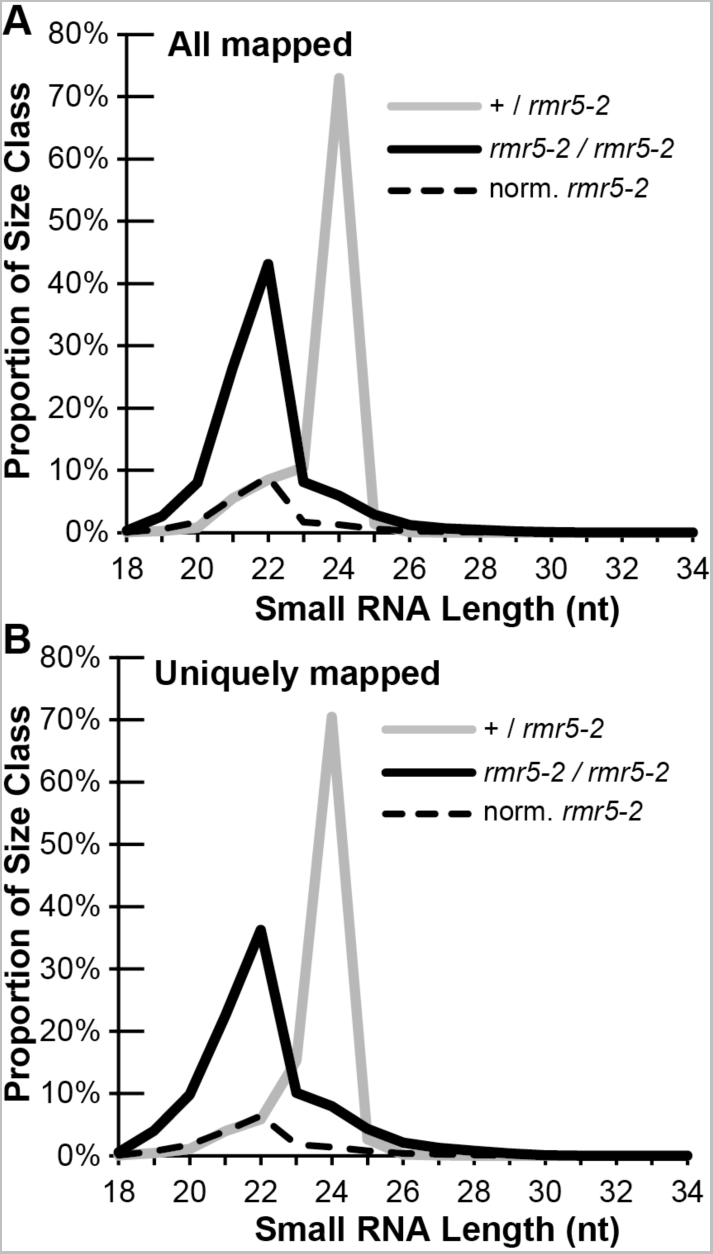
RMR5-Dependent sRNAs. **(A)** Line plot comparing proportions of total genome-matched sRNA sequences versus their length in nucleotides. Both total proportions for the indicated sibling genotypes (black = mutant, grey = non-mutant) as well as normalized proportions of mutant sRNAs (dashed black) relative to the levels of non-mutant 22-nt species are plotted. **(B)** Line plot similar to that in **(A)** comparing proportions of unique, genome-matched, sRNA sequences.

### ZmDCL3 Represses *Pl****′*** Transcription

In inner husk leaves, transcriptional repression of *Pl′* relative to *Pl-Rh* (Hollick et al., 2000) is dependent on both RPD1 (Hollick et al., 2005) and RMR2 (Barbour et al., 2012), whereas RMR1 confers *Pl′* mRNA instability but does not repress transcription (Hale et al., 2007). Because functional *pl1* alleles also confer pigmentation to seedling leaf sheaths, we tested whether the pigment increase seen in *dcl3* mutants was correlated with increased *Pl1-Rh* transcription using a nuclear run-on assay. Radiolabeled nascent RNA generated from isolated nuclei (*dcl3-2* mutant and homozygous non-mutant F_2_ seedlings at 14 days post-imbibition) was hybridized with immobilized riboprobes representing *ubiquitin2*, *pl1*, and *anthocyaninless1* (*a1*), a gene whose transcription is dependent on PL1 (Cone et al., 1993). Relative to *ubiquitin2*, we found significantly higher *pl1* and *a1* transcription rates in mutants (Figure 3). The approximately 4-fold increase in *Pl1-Rh* transcription rate mirrors that seen in *rpd1-1* mutants (Hollick et al., 2005) and is similar to the 3-fold difference observed between *Pl-Rh* and *Pl′* (Hollick et al., 2000). Although it remains unknown if mRNA stabilities are also affected, these results show that ZmDCL3 represses *Pl1-Rh* transcription associated with the paramutant *Pl′* state.

**Figure 3.**
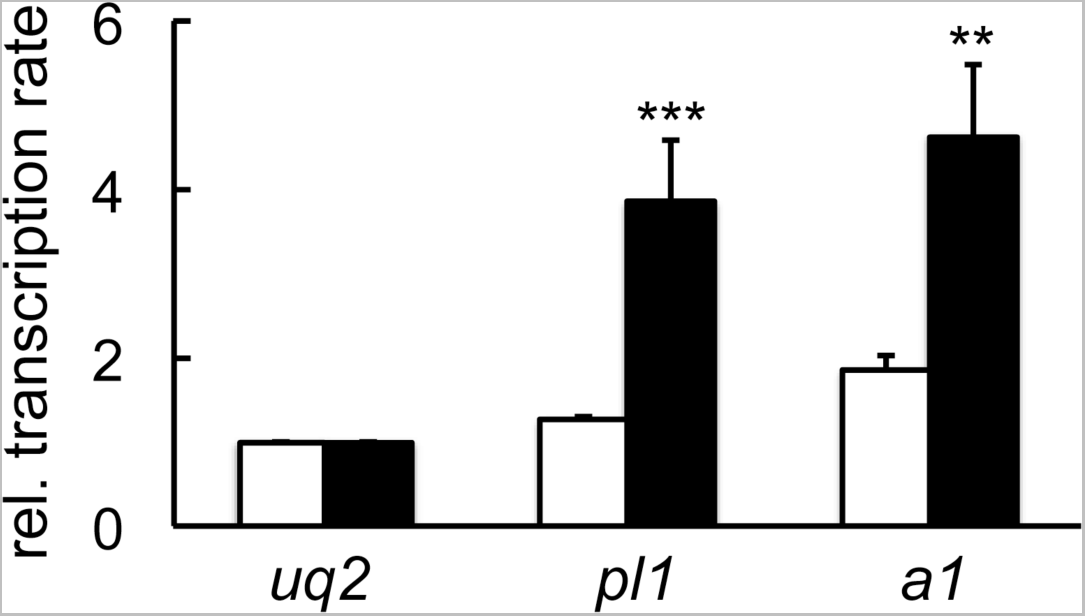
ZmDCL3 Represses *Pl′* Transcription. Histograms represent mean transcription rates (+ / − s.e.m.) of *Pl1-Rhoades* (*pl1*) and *anthocyaninless1* (*a1*) relative to that of *ubiquitin2* (*uq2*) in homozygous non-mutants (white bars) and *dcl3-2* F_2_ sibling seedlings (black bars; n=5 for each). Means were compared using a two-tailed Student’s *t*-test; * -*p* = 0.013; ** - *p* = 0.007.

### Cytosine Methylation is Largely Unaffected in *dcl3* Mutants

Most maize genes exist between so-called mCHH islands (Gent et al., 2013) where RNAP IV / MOP1 generate 24-nt RNAs (Gent et al., 2014; Li et al., 2014; Erhard et al., 2015) that are thought to maintain these 5mC patterns via an RdDM-like mechanism (Gent et al., 2014; Li et al., 2014). Because transcription from *Pl′* states is greater in *dcl3* mutants, we explored possible correlations with ZmDCL3-dependent 5mC at the *Pl1-Rh* promoter-proximal CACTA-type *doppia* DNA transposon fragment, a region where some 5mC patterns are dependent on RMR1 (Hale et al., 2007), MOP1 (Hale et al., 2007), and RPD1 (Hale et al., 2007, Erhard et al., 2013). Primers flanking the *doppia* fragment were used to PCR amplify intervening sequences from bisulfite-treated genomic DNA isolated from terminal flag leaves of *dcl3-2* / *dcl3-2* and + / *dcl3-2* siblings and individual amplicons were then cloned into plasmid vectors and sequenced. We found an extraordinarily high number of 5mC residues within the *doppia* fragment in all samples (Figure 4A) while the immediate flanking sequences had relatively little 5mC (Figure 4B), even though mCHH levels (∼20%) were still above the ∼5% genome average (Regulski et al., 2013; Gent et al., 2014; Li et al., 2014). Approximately 92% of all the *doppia* fragment cytosines, including 86% of those in CHH context, were methylated (Figure 4C) in non-mutants. Overall, ZmDCL3 absence had a modest effect on 5mC levels across these sequences (Figures 4B, 4C and Figure 5A). Only mCG and mCHG levels were significantly lower across the *doppia* fragment (2-sample *t*-test; 0.99 vs 0.96, *t* = 2.61, 30 d.o.f., *p* = 0.014; 0.93 vs 0.83, *t* = 2.83, 30 d.o.f., *p* = 0.008). By confining our analysis to just those 5′ boarder sequences evaluated previously in *rpd1 / rmr6* mutants (Erhard et al., 2013), we found significant differences only in mCG levels (Supplemental Figure 2). These data emphasize that the proximal *doppia* 5mC patterns are largely maintained independent of ZmDCL3 function; a finding in contrast to the RMR1/MOP1/RPD1-dependent mCHH detected within the *Pl1-Rh doppia* fragment using the *Stu*I methylation-sensitive restriction enzyme (Hale et al., 2007) and the MOP1/RPD1- dependent patterns typical of flanking mCHH islands (Li et al., 2014).

**Figure 4.**
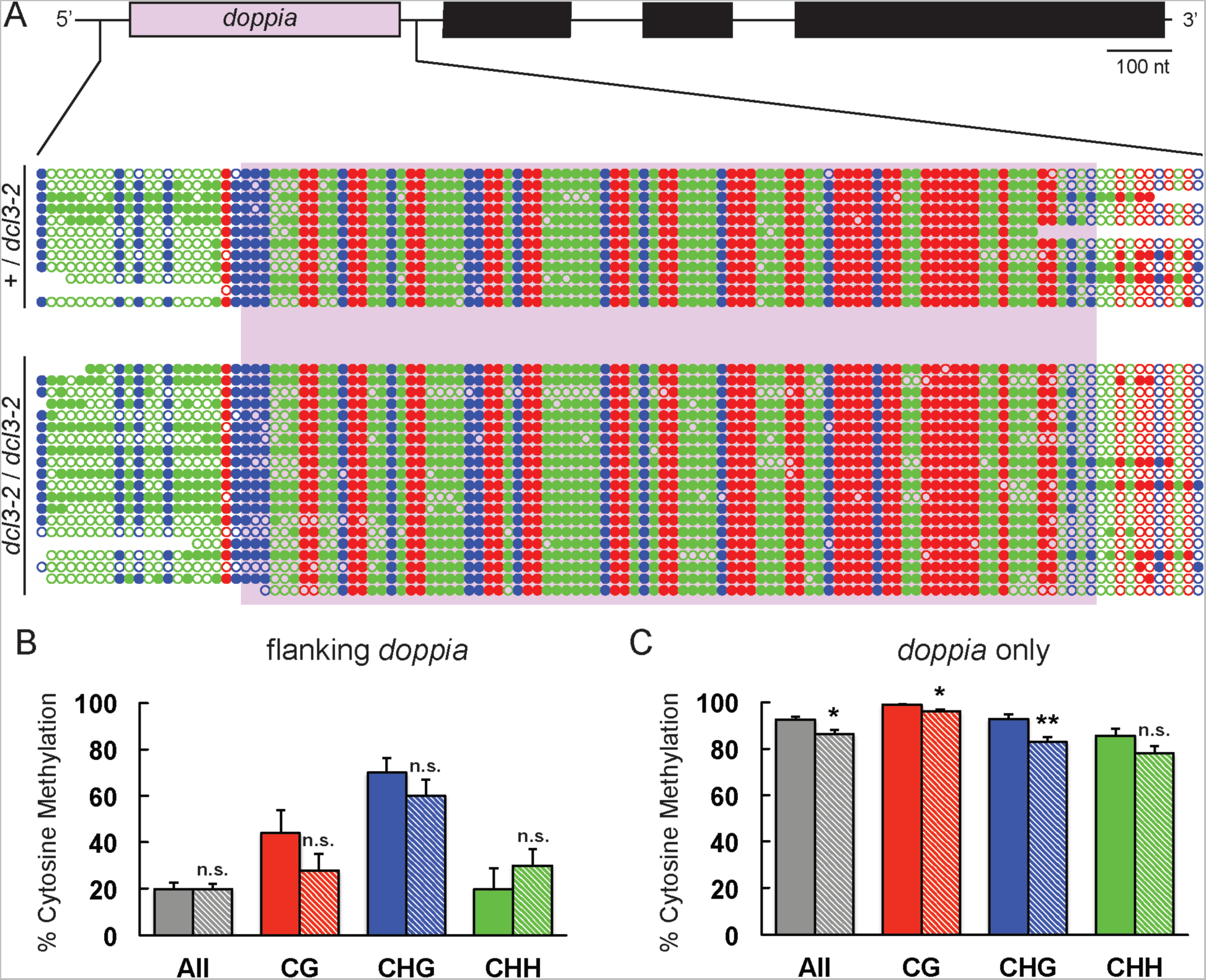
ZmDCL3-Dependent Cytosine Methylation at *Pl1-Rhoades*. **(A)** A schematic of *Pl1-Rhoades* with 5′ *doppia* fragment highlighted in lavender and corresponding dot plots showing methylation status (open = C, solid = 5mC) of each cytosine. Red, blue, and green dots correspond to CG, CHG, and CHH contexts, respectively. **(B and C)** Bar plots representing cytosine methylation levels (means ± s.e.m.) are compared between *+* / *dcl3-2* (solid bars) and *dcl3-2* / *dcl3-2* (hatched bars) genotypes across the flanking regions **(B)** and *doppia* region **(C)** for each cytosine context. Total cytosine methylation levels are indicated by grey bars. Means were compared using a two-tailed Student’s *t-*test; * -*p* < 0.02, ** -*p* < 0.006.

**Figure 5.**
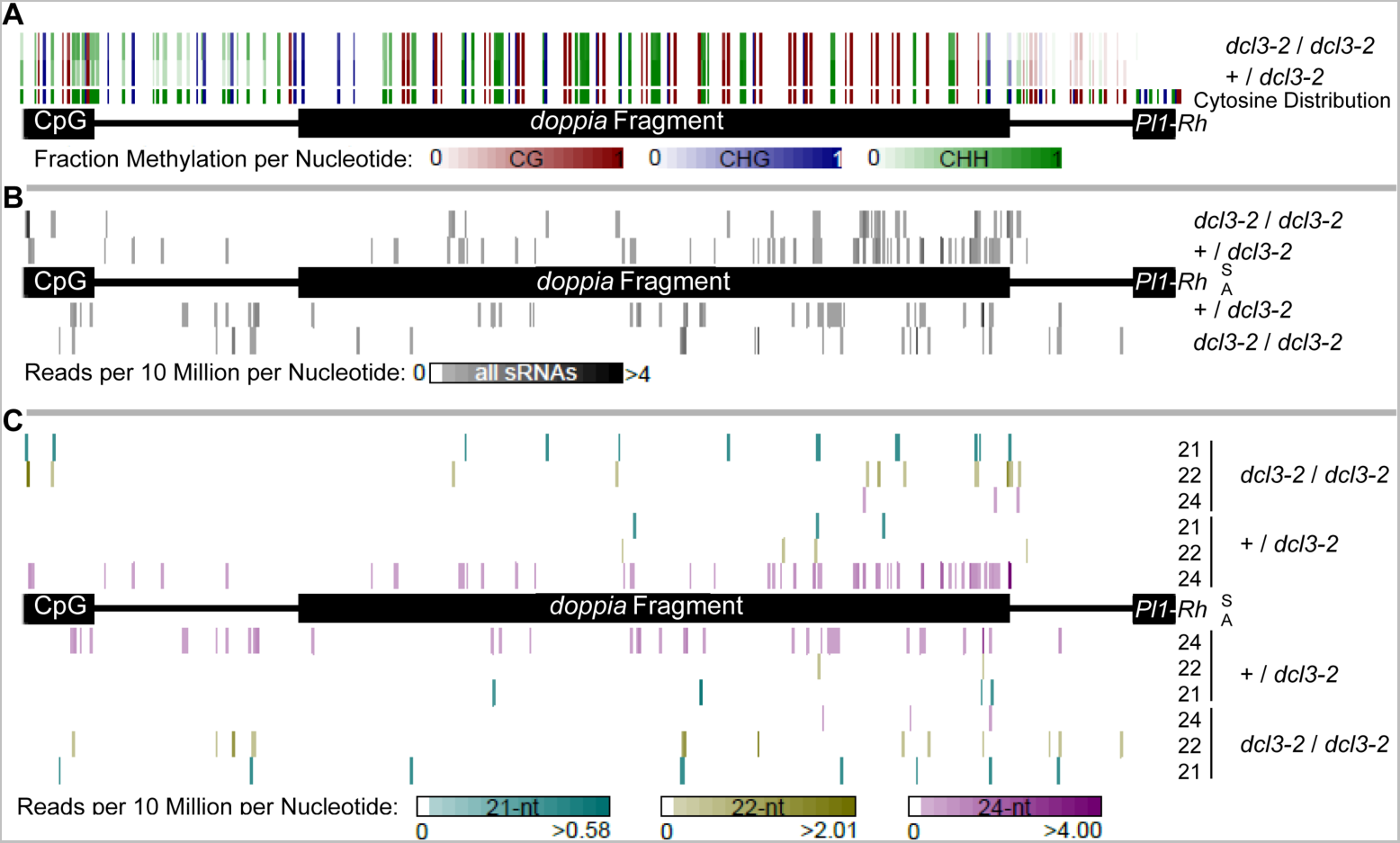
ZmDCL3-Dependent sRNAs Mapping to the *Pl1-Rhoades* Promoter. **(A)** Fractions of cytosines methylated per context across the bisulfite-sequenced *Pl1*-*Rhoades* region are displayed for the indicated genotypes. Red, blue, and green correspond to CG, CHG, and CHH contexts respectively. **(A-C)** Black boxes highlight an upstream CpG-rich region (Gross 2007), *doppia* fragment, and first exon of the *Pl1-Rhoades* gene model. **(B)** Total sRNA abundances mapping across the bisulfite-sequenced *Pl1*-*Rhoades* region are displayed for the indicated genotypes. Sense (S) and antisense (A) alignments are shown and only the most 5′ nucleotide of a given read is indicated. **(C)** Abundances for 21-, 22-, and 24-nt RNAs at the bisulfite-sequenced *Pl1-Rhoades* region are given for the indicated genotypes. Sense (S) and antisense (A) alignments are shown and only the most 5′ nucleotide of a given read is indicated. Cyan, yellow, and magenta correspond to 21-, 22-, and 24-nt small RNAs, respectively.

We also used a chop-PCR assay to test the ZmDCL3 requirement in maintaining 5mC patterns at other known RPD1 targets including *CRM* and *Prem2* LTR-retrotransposons (Hale et al., 2009) and a specific LTR-gypsy element of the *ubid* subclass found upstream of the *outer cell layer 2* (*ocl2*) gene (Erhard et al., 2015). A series of enzymes with collective sensitivities to methylated cytosines in all sequence contexts were used and the results (Supplemental Figure 3) indicated no discernable differences in enzyme digestion patterns due to ZmDCL3 absence. Even a methylated chromosome *1* satellite repeat had no ZmDCL3-related differences (Supplemental Figure 3).

### ZmDCL3 Absence Alters *doppia* sRNA Profiles

In Arabidopsis *dcl3* mutants, both DCL2 and 4 provide different-sized sRNAs that may serve as RdDM-type effectors (Xie et al., 2004; Gasciolli et al., 2005; Henderson et al., 2006; Kasschau et al., 2007; Stroud et al., 2013). Even DCL-independent RNAP IV sRNAs that are exonuclease trimmed to effective sizes are sufficient to maintain a subset of 5mC patterns (Ye et al., 2016; Yang et al., 2016). We therefore hypothesized that the 5mC maintenance within the *doppia* fragment might be similarly due to either functional DCL redundancy or DCL-independent sRNAs. Focusing on the immediate 5′ region of *Pl1-Rh*, we aligned all 21, 22, and 24-nt reads from the immature cob sRNA libraries and compared their normalized abundances (Figure 5). Abundances and distributions of these sRNA reads were fairly conserved in both *dcl3-2* mutants and their non-mutant siblings (Figure 5B) but the 24-nt class was largely ZmDCL3 dependent (Figure 5C). The sRNA reads mapping adjacent to the *doppia* fragment in both mutant and non-mutant genotypes indicate the *Pl1-Rh* 5′ promoter region is transcribed by RNAPs in immature cobs.

### Alternate Sized sRNAs Replace 24mers in the Absence of ZmDCL3

Because sRNAs sampled from *dcl3* mutants represent a fraction depleted of most 24mers (approximately 65% of total normal sRNA levels), the relative abundance of other sRNA sizes is greatly increased. We attempted to address whether ZmDCL3 substrates are processed into other size classes in *dcl3* mutants by defining 23/24mer-generating clusters over the first 100 Mb of chromosome *1* and then asking whether alternative sized sRNAs appeared in *dcl3-2* mutants (see Supplemental Methods and Supplemental Dataset 1). We identified 6,099 total clusters, of which 1,796 were exclusively occupied by 23/24mers. Nearly half (819) of these exclusive clusters showed greatly reduced sRNAs without ZmDCL3 (fraction of hits-normalized 20-25mer sRNAs in *dcl3-2 / dcl3-2* compared to sibling *+ / dcl3-2* were less than 25%) with no sRNAs found in the majority (478). Thus, in the absence of ZmDCL3, nearly 1/3 of all ZmDCL3- dependent sRNA generating loci either do not produce a dsRNA substrate or such RNAs are not processed by alternative dicer enzymes. Meanwhile, 20% of clusters had similar (75-125%) or increased (>125%) total sRNAs in the *dcl3-2 / dcl3-2* immature cobs, presumably representing ongoing transcription at these loci. Reviewing sRNA abundances at these transcriptionally-active clusters, most (348 of 361, 96.4%) had tallies for multiple sRNA sizes. 22mers accounted for over half of the sRNAs from 183 of the 361 active clusters (50.7%). Overall, these data indicate that multiple dicing or degradation pathways exist for most cluster-originating RNA transcripts produced in the absence of ZmDCL3.

### ZmDCL3 Does Not Affect *ocl2* mRNA Levels

In the absence of RPD1, transcription rates are increased at both the *ocl2* gene and its 5′ - proximal *ubid_RLG* element (Erhard et al., 2015). Because *ocl2* mRNA levels were similarly increased (Erhard et al., 2015), we asked whether they were also elevated in *dcl3* mutant seedlings. Quantitative RT-PCR amplifications of *ocl2* cDNA normalized to *alanine aminotransferase* (*aat*) levels showed no statistical differences between *dcl3-2* mutants and non-mutant siblings (Figure 6) in contrast to the significant 10.9-fold elevations (two-tailed Student’s *t*-test, *p* = 0.011) seen in the absence of RPD1. This result represents the third documented example in which RPD1 regulatory functions are separable from the production of 24-nt RNAs (Hale et al., 2009), and the first case of an allele regulated by RNAP IV functions independent of 24-nt effectors.

**Figure 6.**
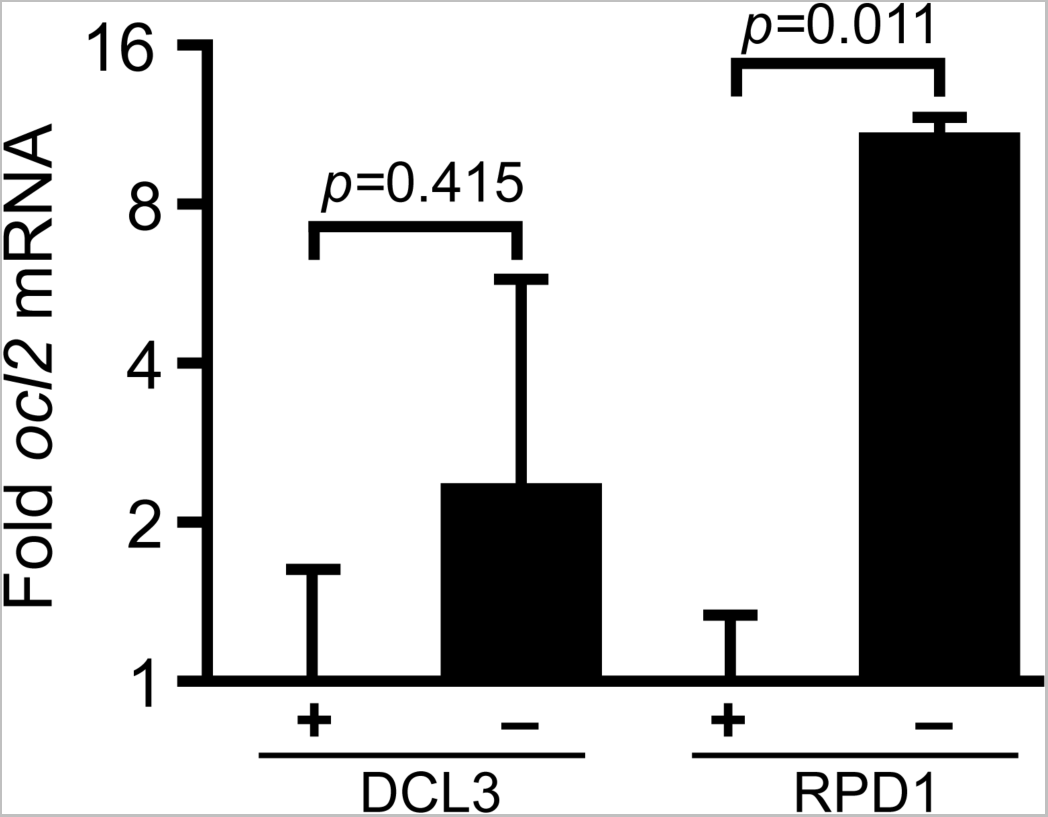
RPD1-Specific Effects on *ocl2* mRNA Levels. Mean fold *ocl2* RNA (2^-ΔΔCt^)(±s.e.m) in three pooled 8-day seedlings homozygous for *dcl3-2* (- / -) mutants or *rpd1-1* (- / -) mutants relative to respective non-mutant siblings (+ / +). The *ocl2* mRNA levels are normalized to *aat* mRNA abundance. Means were compared using a two-tailed Student’s t-test.

### *dcl3* Mutations Affect Plant Development

In contrast to RPD1 defects (Parkinson et al., 2007; Erhard et al., 2009), the dark pigment *Pl-Rh*-like phenotypes observed in homozygous *dcl3* mutant plants appear unassociated with severe developmental abnormalities. However, minor changes in both plant heights and flowering times correlated with ZmDCL3 function (Figure 7A, 7B) as assessed from field-grown progenies segregating 1:1 for *dcl3-2* (121283 and 121279) and *dcl3-4* (120672) mutants and their respective heterozygous siblings. Plant heights were significantly affected by the absence of ZmDCL3 in both the first (May 27) and second (June 6) plantings. Curiously, the *dcl3-2* mutants were taller than the respective non-mutant siblings in the May 27 planting (121283), in contrast to the reduced plant heights observed in the other three plots. Despite this unexplained difference in plant heights, flowering times were significantly delayed in *dcl3* mutants in all but one plot – May 27 planting of the 120672 progeny. Similar trends of both plant height and flowering time differences were found in previous smaller plantings of both 121279 (*dcl3-2*) and 120672 (*dcl3-4*) F_2_ progenies (see Supplemental Figure 4) but only the *dcl3-4* mutants showed consistently significant differences. These correlations with independent mutant *dcl3* alleles in different genetic backgrounds (121283 and 121279 – 60% A632 and 27% A619; 120672 – 95% A619) provide evidence that ZmDCL3 plays a role in ensuring normal growth and development.

**Figure 7.**
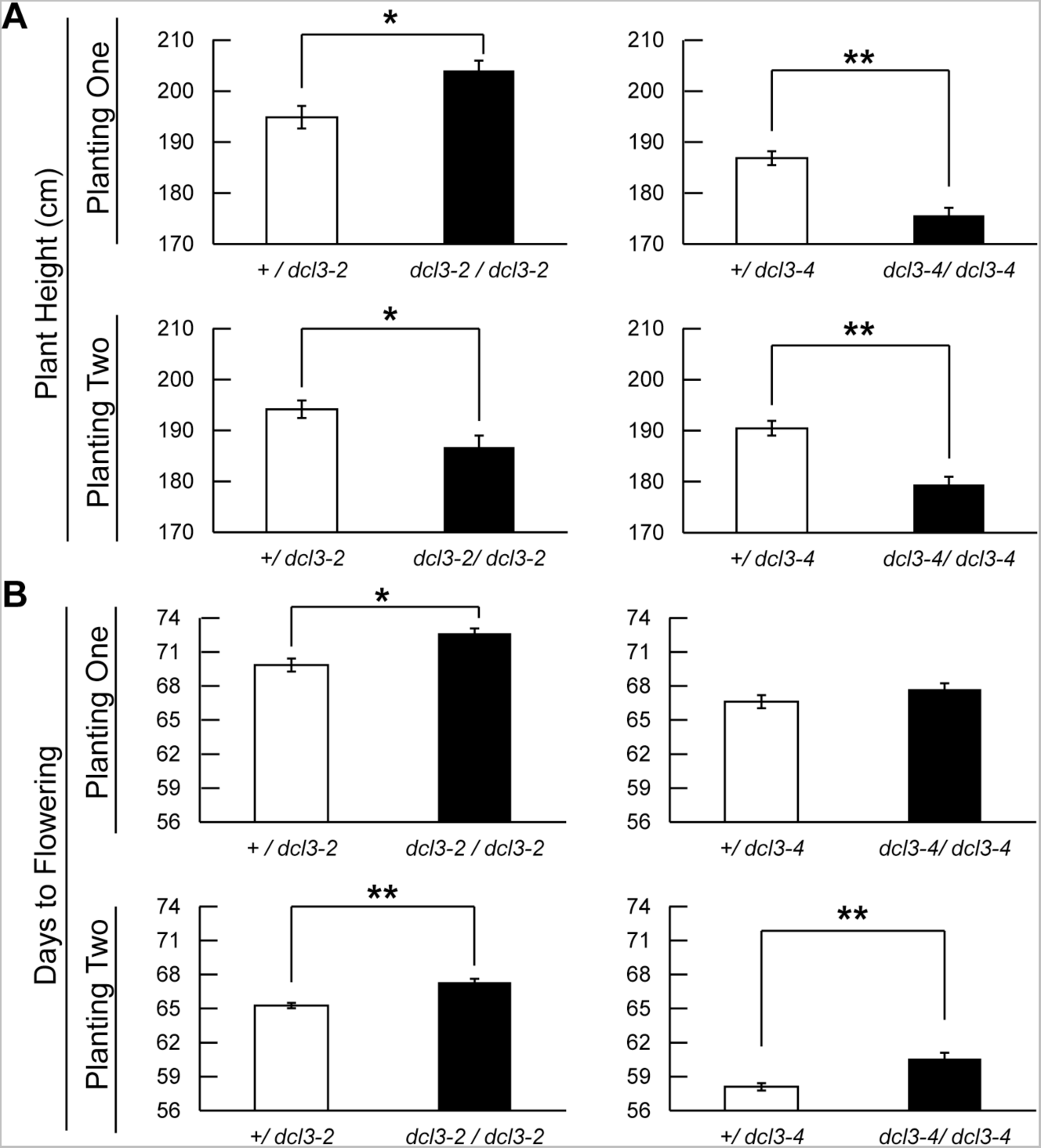
ZmDCL3 Effects on Plant Development Bar plots represent. **(A)** mean plant heights (+ / − s.e.m.) and **(B)** mean days to flowering (+ / − s.e.m.) for sibling plants of the indicated genotypes planted two weeks apart in 2014. Sibling means were compared using a two-tailed Student’s *t-*test; *- *p* < 0.01; **- *p* < 5.0 x 10^-4^. Planting 1: + / *dcl3-2* (n = 40); *dcl3-2* / *dcl3-2* (n = 41); *+* / *dcl3-4* (n = 45); *dcl3-4* / *dcl3-4* (n = 50). Planting 2: + / *dcl3-2* (n = 75); *dcl3-2* / *dcl3-2* (n = 57); + / *dcl3-4* (n = 51); *dcl3-4* / *dcl3-4* (n = 34).

## Discussion

In the developing maize female inflorescence – and presumably all vegetative tissues – the sole dicer-like3 protein generates 24-nt RNAs from both repetitive and unique genomic origins. The related grass-specific ZmDCL5, in contrast, is specialized to produce phased, stamen-specific 24-nt RNAs (Song et al., 2012) from 176 dispersed loci (Zhai et al., 2015b) affecting male fertility (Teng et al, 2020). *In vitro*, DCL3 preferentially binds short dsRNAs (Nagano et al., 2014) with 3′ overhangs and coordinates the precise cleavage of either 24-nt/24-nt or 24-nt/23-nt duplexes using its tandem RNase III domains (Loffer et al., 2022). Three of the four *dcl3* alleles identified in our mutant screen have early premature stop codons that would encode peptides lacking both RNase III domains. Additionally, the *dcl3-2* nonsense mutation occurring within the N-terminal DEAD-box helicase domain-coding region should abolish any partial or inhibitory functions from any truncated peptides. Consistently, ∼95% of all 24mers from 4.5cm immature cobs are absent in the presumed genetic null *dcl3-2* homozygotes.

### ZmDCL3 non-redundancy

The fact that *dcl3* was identified in our genetic screen for *rmr* loci indicates that there is no functional redundancy of other maize dicer-like proteins for maintaining *Pl′* repression. Overall, our analyses indicate that relatively few sRNA precursors are processed to alternative sizes, and that these are insufficient to specifically maintain *Pl′* transcriptional repression and induce *b1* paramutations, and more generally to control normal growth and reproductive timing. Other examples of non-redundant DCL3 functions are found in *Physcomitrella patens* (moss) (Cho et al., 2008) where *Ppdcl3* mutants have an increased rate of gametophore production; in rice where inverted-repeat *Osdcl3* lines result in dwarfism, smaller panicles, and increased leaf blade angle (Wei et al., 2014); and in *Nicotiana atteunata* where an inverted-repeat *Nadcl3* line increases leaf length and width (Bozorov et al., 2012). The DCL redundancy seen in Arabidopsis (Xie et al., 2005; Gascolli et al, 2005; Henderson et al., 2006; Kasschau et al., 2007; Stroud et al., 2013) may extend to *Solanum lycopersicum* (tomato) and *Nicotiana benthamiana* in which DCL3 RNAi-based knockdowns have no effect on plant growth and development (Kravchik et al., 2014; Katarou et al., 2019). Surprisingly, we found ZmDCL3 was not required to maintain the repressed *B′* expression state, unlike MOP1 (Dorweiler et al., 2000), RPD1 (Hollick et al., 2005) and RP(D/E)2a (Stonaker et al., 2009; Sidorenko et al., 2009). This finding implicates either functional redundancy provided by other ZmDCL proteins or a direct requirement for normal RNAP IVa-dependent dsRNA production at the *B1-I* allele.

In contrast to the ZmDCL3-independent maintenance of repressed *B′*, we found ZmDCL3 essential to facilitating paramutation in *B′* / *B-I* heterozygotes. In accord with an RdDM-type model, this induction requires the 24-nt RNA biogenesis carried out by RNAP IVa (Hollick et al., 2005; Stonaker et al., 2009; Sidorenko et al., 2009), MOP1 (Dorweiler et al., 2000), and ZmDCL3. Although, an interesting exception occurs when the *B′* -contributing female parent is homozygous for the *rp(d/e)2a-1* mutation (Stonaker et al., 2009). Thus, 24-nt RNAs are one potential source of paramutagenic function that might account for the *trans*-homolog communication typifying paramutation-type interactions (Alleman et al., 2006; Erhard et al., 2009; Stonaker et al., 2009; Sidorenko et al., 2009; Arteagua-Vasquez et al., 2010; Belele et al., 2012; Greaves et al., 2012; Gouli et al., 2016). Whether ZmDCL3 is similarly required for mediating *pl1* paramutation remains to be adequately evaluated as these tests are complicated by the reversions of *Pl′* to *Pl-Rh* occurring in *dcl3* mutants.

For most examples of paramutation described so far in maize, a transcriptionally-repressed state is always associated with robust paramutagenic activity persisting through meiosis (Patterson et al., 1993; Hollick et al., 2000; Sidorenko and Chandler 2008). Derepression occurring in various *rmr* or *mop* mutant backgrounds, however, does not always mitigate or eliminate the meiotic inheritance of paramutagenic action. This indicates that, to some extent, the mechanism(s) responsible for transcriptional repression are distinct from those maintaining the meiotically-heritable feature responsible for paramutagenic function. This idea is further supported by the progressive loss of *Pl1-Rh* paramutagenicity seen following *trans*-generational absence of RMR1 (Barbour et al., 2012), RP(D/E)2a (Stonaker et al., 2009), and ZmDCL3. Thus, while high gene expression levels in the previous generation appear necessary for losses of paramutagenic activity, it is often insufficient. Indeed, *B′* always remains paramutagenic following transmission from *mop1*, *rp(d/e)2a* /*rmr7*/*mop2*, and *rpd1*/*rmr6* mutants (Dorweiler et al., 2000; Stonaker et al., 2009; Sidorenko et al., 2009; Hollick et al., 2005), and even *Pl′* remains paramutagenic following transcriptional reactivation in *rmr2-1* mutants and a subsequent seven generations of self-pollinations (Barbour et al., 2012). Thus, the mechanism(s) that maintain paramutagenicity in certain *rmr* mutants appear unable to compensate for persistent loss of sRNA biogenesis over multiple generations indicating that this mechanism requires occasional reinvigoration.

### Maintenance of 5mC patterns

Most maize genes are flanked by RNAP IV function (Gent et al., 2013; Li et al., 2014; Li et al., 2015b; Erhard et al., 2015) that enhance RNAP II terminations (Erhard et al., 2015). These flanking regions are coincidently enriched for 24-nt RNAs and mCHH, but relatively depleted of mCHG and mCG (Gent et al., 2013; Li et al., 2014; West et al., 2014). In Arabidopsis, mCHH can be maintained independent of RdDM via CMT2 bound to H3K9me2 (Du et al., 2012; Du et al, 2014; Stroud et al., 2014). Because maize lacks a CMT2-type protein (Zemach et al., 2013), and global mCHH levels are reduced in *mop1-1*, *mop2*, and *mop3-1* mutants (Li et al., 2014; Li et al., 2015b), it is inferred that most maize mCHH is dependent on an RdDM-type mechanism. Because Arabidopsis RdDM facilitates *de novo* 5mC in all sequence contexts (Cao and Jacobsen 2002), it is unclear why these RNAP IV-engaged regions flanking most maize genes are largely devoid of mCHG and mCG. The CACTA-type *doppia* fragment immediately upstream the *Pl1-Rh* transcription unit is highly unusual in having nearly every cytosine fully methylated in flag leaves. More remarkable is our finding that this pattern persists in the absence of ZmDCL3, even in this fully-differentiated terminal tissue. We consider several possibilities to account for this observation and discuss these below. Either these patterns are faithfully maintained via 1) maintenance methylation, 2) RdDM-independent *de novo* methylation, 3) DCL/sRNA redundancy, 4) demethylation inhibition, or some combination(s) thereof.

#### Maintenance methylation

In Arabidopsis, mCG and mCHG patterns are largely maintained by the DMNT1-like METHYLTRASFERASE 1 (MET1) and CMT3, respectively (Kishimoto et al., 2001; Lindroth et al., 2001; Cao and Jacobsen, 2002; Cao et al., 2003). CMT3 can also contribute to mCHH maintenance (Bartee et al., 2001; Lindroth et al., 2001; Cao and Jacobsen, 2002; Cao et al., 2003; Stroud et al., 2013). The two maize CMT3-type orthologs encoded by the *zmet2* and *zmet5* genes (Papa et al, 2001; Makarevitch et al., 2007; McCarty and Meeley 2009) are similarly required for some mCHH, particulary in small Type II DNA transposons (>1 kb) (Li et al., 2014). Although it’s formally possible that all the *doppia* mCHH is reflective of maintenance methyltransferases, we know of no precedence for such extensive activity. Moreover, in the absence of any possible *de novo* methylation, these patterns would need to be parentally inherited. In contrast, both mCG and mCHG levels across this *doppia* fragment are slightly decreased in the *dcl3-2* mutant, consistent with a role for ZmDCL3-independent RdDM acting on these specific regions at some point in ontogeny.

#### RdDM-independent de novo methylation

In rice, OsDCL3 produces 24-nt centromeric OsCentO siRNAs, MITE-derived siRNAs and noncanonical long microRNAs (Wu et al., 2010; Yan et al., 2011; Song et al., 2012) including CACTA-TE-derived mir820 that targets the major *de novo* cytosine methyltransferase-encoding gene, *OsDOMAIN REARRANGED METHYLTRASFERASE 2* (*OsDRM2*) (Wu et al., 2010, Nosaka et al., 2012, Moritoh et al., 2012). Like the *Pl1-Rh doppia* fragment, most rice MITEs appear to retain mCG, mCHG, and mCHH modifications and it is proposed that elevated OsDRM2 might counteract the loss of 24-nt-based targeting (Wei et al., 2014), but how this might occur remains unaccounted for. It is also unclear whether the maize DRM1/2 orthologs encoded by *zmet3* and *zmet7* might be similarly regulated.

#### DCL/sRNA redundancy

Even though our analyses of 4.5cm cob sRNA profiles show modest replacement of 24mers with other size classes, it is possible that a non-canonical RdDM-type mechanism – like that proposed to initiate RdDM via RNAP II-derived 21 and 22nt RNAs (Sigman et al., 2021) – remains active in distal leaf blade primordia by using different sRNA size classes. There might also be a more substantial production of these other size classes in these tissues or the relatively few alternative size classes may be sufficient. Thus, there might be functional DCL redundancy in some tissues and not others.

#### Demethylation inhibition

In Arabidopsis, active cytosine demethylation is carried out by DEMETER-LIKE (DML) DNA glycosylases and subsequent base-excision repair (Gong et al., 2002; Gehring et al., 2006; Ortega-Galisteo et al., 2008). Thus, existing 5mC patterns are the result of initial *de novo* methylation, subsequent maintenance methylation, and active demethylation events (reviewed in Law and Jacobsen 2010, and in Zhang et al., 2018). There appear to be multiple mechanisms for DML recruitment by either specific histone H3 covalent modifications (Qian et al., 2012; Tang et al., 2016) and/or a large protein complex containing a 5mC-binding protein (Lang et al., 2015; Duan et al., 2017). It is possible that maize DML orthologs are prohibited from acting on this *Pl1-Rh doppia* fragment and other repetitive sequences we profiled.

### Regulatory role(s) of the *Pl1-Rh doppia* sequences

How the *Pl1-Rh doppia* fragment affects paramutation and/or gene regulation remains unclear. A very similar *doppia* fragment constitutes over half of the entire 5′ sequence found between divergently oriented *r1* genes (Walker et al., 1995) that are aleurone expressed (Stadler 1946). The *Pl1-Blotched* allele, which is nearly identical to *Pl1-Rh* over the compared sequences (Hoekenga 1998), is typically aleurone-expressed in a variegated pattern (Cocciolone and Cone, 1993) but achieves uniform pigmentation in the absence of RPD1 (Erhard et al., 2013). Even *Pl1-Rh*, which is not typically aleurone expressed, can be conditioned to produce full aleurone pigmentation through repeated loss of RPD1 function (Erhard et al., 2013). These results indicate that RNAP IV normally represses aleurone-specific regulatory sequences contained within the *doppia* fragment. Aleurone pigmentation was also associated with *mop1*, *rmr1*, and *rp(d/e)2a*/*rmr7* mutant kernels consistent with an RdDM-like repression mechanism (Erhard et al., 2013).

Despite the presence of an identical *doppia* fragment, *Pl1-Blotched* is not rendered paramutagenic following exposure to *Pl′* (Hollick et al., 2000; Gross and Hollick 2007). This indicates that the *doppia* sequences by themselves are insufficient for *Pl1-Rh* paramutation behaviors. Furthermore, a recombinant allele (*pl1-R30*) between the *Pl1-Rh* coding region and *pl1-B73* 3′ sequences confers weak, or no, pigmentation to various plant parts and, like *Pl1-Blotched*, cannot acquire paramutagenic function (Erhard et al., 2013). Thus, while the *doppia* sequences may be necessary for paramutation, their presence is insufficient. These observations point to sequences >6kb 3′ of the *Pl1-Rh* coding region that are essential for both high transcription levels and for facilitating paramutation (Erhard et al., 2013). Perhaps these important regulatory sequences are also targeted by an RdDM-type mechanism where ZmDCL3 plays a non-redundant role in specifying chromatin modifications.

### RdDM-independent roles for maize RNAP IV

Outside of its role in paramutation, maize RPD1 has been co-opted to define tissue- and/or development-specific expression for many genes, including *silkless1* (Parkinson et al., 2007) and *ocl2* (Erhard et al., 2015). Correspondingly, *rpd1*/*rmr6* mutants display defects in tassel morphology, floral identity, internode elongation, leaf polarity, delayed phase change, and stochastic ectopic outgrowths (Parkinson et al., 2007; Erhard et al., 2009). These defects appear to reflect a 24-nt RNA-independent role for RNAP IV because similar developmental defects are not seen *mop1*, *rmr1*, *rmr2*, or *dcl3* mutants which all lack significant 24-nt RNA biogenesis (Nobuta et al., 2008; Hale et al., 2007; Barbour et al., 2012). Additionally, RPD1 absence facilitates accumulation of RNAP II transcripts from *Prem2/Ji* and *CRM* LTR elements whereas loss of MOP1 or RMR1 does not (Hale et al., 2009). In this study, we found that the 5mC status of these specific LTR TEs and of the *gypsy-ubid* element found upstream of the *ocl2* gene remained unchanged in *dcl3* mutants. In comparison to the increased *ocl2* RNAP II nascent transcription and mRNA abundance found in the absence of RPD1 (Erhard et al., 2015) in day-8 seedlings, we found no increased *ocl2* mRNA accumulation in the absence of ZmDCL3. This finding, along with previous results (Hale et al., 2009), supports the concept that RNAP IV has regulatory functions independent of an RdDM-type mechanism, perhaps through direct competition with RNAP II for genomic templates (Erhard et al., 2015).

Both the expansion and functional specialization of Argonaute-type proteins in *Caenorhabditis elegans* has facilitated diverse biological roles for sRNAs (reviewed in Youngman and Claycomb, 2014) including their direct use for the inheritance of gene regulatory responses to environmental stresses (Rechavi et al., 2014) and dsRNA-induced RNA interference (RNAi) (Fire et al., 1998; Vastenhouw et al., 2006). The inheritance of paramutant regulatory states, at least at *Pl1-Rh*, is similarly dependent on sRNAs but whether this requires transmission of specific sRNAs remains unknown. The common requirement of MOP1 (Dorweiler et al., 2000), RNAP IVa (Hollick et al., 2005, Stonaker et al, 2009; Sidorenko et al., 2009), and ZmDCL3 to facilitate paramutation at *b1* is consistent with an RdDM-type mechanism involving 24-nt RNAs, yet additional details of this model remain to be tested. In maize and other grass species, the retained duplications of genes encoding both RNAP IV catalytic subunits and DCL3-type proteins provide a particularly interesting opportunity for functional diversification of transcriptional control and sRNA-mediated biology. In rice, RNAi-based knock-down of the duplicate *OsNRPD1* genes leads to either no effects (Debladis et al., 2020) or increased tillers and shorter stature (Zhang et al., 2020), and *OsDCL3* knock-down affects the regulation of *OsDRM2* and brassinosteroid biosynthesis genes (Wei et al., 2014). In maize, *ocl1* mRNA translation is regulated by a MOP1 / RPD1-dependent 24-nt RNA (small1) having complementary to the *ocl1* 3′ UTR (Klein-Cosson et al., 2015). Thus, breeding efforts for the two most globally important crop species has led to alternative co-opted uses of sRNAs and core RdDM-type components, some for agricultural success.

## METHODS

### Genetic Materials and Stock Syntheses

The genetic nomenclature used follows established guidelines for maize (*Zea mays*) and has been fully described previously (Hollick et al., 2005). Female-derived factors are placed before male-derived factors when stating diploid genotypes. Anther pigmentation is visually assessed using a 1 – 7 graded Anther Color Score (ACS) as previously described (Hollick et al., 1995).

Hand pollinations were used for all stock syntheses and genetic assays. Detailed pedigree information is available upon request. All color factor alleles including *B1-Intense* (*B1-I*) and *Pl1-Rhoades* (*Pl1-Rh*) used herein derive from a color-converted W23 line developed by Ed Coe, Jr. (USDA-ARS, University of Missouri, Columbia, MO) and maintained by selfing and sib crosses since 1994. Spontaneous *B′*, *Pl′*, and *B′ Pl′* derivatives from Coe’s original W23 *B-I*, *Pl-Rh* lines have been identified (Hollick 2010) and maintained by selfing. The specific color-converted A619 and A632 lines used here are described in Hollick et al. (2005). All plants are homozygous for the *Pl1-Rh* haplotype unless otherwise described. Stocks used for pollen mutagenesis with ethyl methanesulfonate and subsequent screening criteria for *rmr*-type mutations have been described previously (Hale et al., 2007).

For the *b1* paramutation tests, pollen collected from a single *b1* / *b1* ; *rmr5-1* / *rmr5-1* individual was divided and applied to receptive ears of isogenic *B-I* / *B-I* ; *Pl′* / *Pl′* and *B′* / *B′* ; *Pl-Rh* / *Pl-Rh* plants. Resulting F_1_ individuals were intercrossed and the offspring (progeny identifier 080003) were evaluated for both plant and anther colors. Thirteen of 75 (0.17) individuals had dark anthers expected of *rmr5-1* homozygotes and 20 of 75 (0.27) had no plant color (*b1* / *b1*). All 11 plants having dark anthers and some evidence of plant color were crossed by a recessive *b1* / *b1* tester representing the K55 inbred line and progeny were evaluated for plant color.

For the methylation analysis, sRNA libraries, and phenotypic comparisons, lines were derived from a single homozygous *rmr5-2* M_2_ plant. Individuals from a M_2_S_2_ progeny were backcrossed to a A632 color-converted line (*B-I* / *B-I* ; *Pl′* / *Pl′*) and resulting F_1_ individuals were crossed by related M_2_S_2_ homozygous *rmr5-2* plants to generate progenies segregating 1:1 for homozygous mutants and heterozygous siblings. The 1:1 segregating lines were then maintained by sib crossing. The mutant (*rmr5-2* / *rmr5-2*) and non-mutant (*Rmr5* / *Rmr5*) individuals used for qRT-PCR comparisons were siblings from a BC_1_F_2_ progeny (130009) derived using a B73 *T6-9* line (Hollick et al., 2005) as the recurrent female parent and a single *rmr5-2* mutant from the previously mentioned 1:1 segregating lines. The *rmr5-4* line utilized for the phenotypic comparisons derives from M_2_S_1_ siblings. M_2_S_1_ *rmr5-4* mutants were backcrossed to an A619 color-converted line (*b1* / *b1* ; *Pl′* / *Pl′*) stock with subsequent recurrent F_2_ selection of mutant types and backcrossing to the BC_4_F_2_ generation. Sib crosses of BC_4_F_2_ individuals produced progeny segregating 1:1 for homozygous mutants and heterozygous siblings that was utilized for phenotyping. The segregating 1:1 stocks for *rmr5-2* and *rmr5-4* were planted in four separate plots of 60 kernels each in spring 2013 and again in four separate plots of 120 kernels each in spring 2014. Plant height and days to flowering were recorded for each plant at anthesis (male flowering). BC_1_F_2_ progenies (74% A632) were derived from single *rmr5-1 / rmr5-1*, *rmr5-2 / rmr5-2*, and *rmr5-3* / *rmr5-3* M_2_ individuals for molecular mapping.

### Mapping Primer Design

Oligonucleotide primers for mapping were designed using public SSR marker databases located at MaizeSequence (http://www.maizesequence.org) and MaizeGDB (http://www.maizeGDB.org). Other primers were designed using the PERL script find_ssr (Gross 2007) and Primer3 (http://frodo.wi.mit.edu/primer3). Primers were assayed via PCR with inbred line A619 and A632 genomic DNA samples to determine any polymorphisms present between the two parental inbred lines. Amplicon abundance and size was assayed on 4% agarose gels with ethidium bromide (EtBr) staining. One hundred fifty-one new sets of oligonucleotide primers were designed for use in both bulk segregant and recombination mapping. Relative locations of these primer sets and 78 preexisting pairs found in the public domain are listed in Supplemental Table 3.

### SSLP Mapping

Seedlings were grown under nutrient limitation conditions (sand and tap water only) for 14-16 days and then the uppermost leaf (approx. 0.25 grams) was collected from all putative mutant individuals having dark purple pigmentation in the first seedling leaf sheath. Seedings were then transplanted into pots with top soil, grown to maturity, and anther color scores of sampled individuals were evaluated at flowering to confirm their genotype assignment.

Leaf tissue was lysed in 0.7 mL extraction buffer (1M Tris pH 8.0, 0.5M EDTA, 5M NaCl) using a plastic pestle and electric hand drill. Samples were then treated with 75µL 10% SDS and 17 µL 5M potassium acetate (3M K, 5M Ac), centrifuged at 15,000 rpm for 5 minutes, and supernatants cleared by passing through miracloth (CalBiochem). Genomic DNA (gDNA) was precipitated overnight from the filtered supernatant using 0.7 mL of isopropanol at −20**°**C. gDNA was then pelleted at 15,000 rpm for 3 minutes and resuspended in 100 µL H_2_O.

For bulked segregant analysis, seedling DNA samples from 15 *rmr5-3* mutants were pooled and diluted 1:20 with H_2_O. PCR amplifications from this pool, along with those from A619, A632 gDNAs and synthetic A619/A632 heterozygote controls were produced with the collection of our SSLP markers and visualized on 4% agarose gels. Polymorphisms identified between A619 and A632 control lines were used to determine the approximate genotype of the individuals in our bulked samples. A bias in PCR amplification towards the mutagenized A619 inbred line in the bulked samples was considered evidence for linkage.

For fine-scale recombination mapping, gDNA from a total of 98 *rmr5-1* and *rmr5-2* mutant individuals was tested against additional chromosome *3L* SSLP markers. The number of recombinant chromosomes, representing those individuals with both A619 and A632 genotypes, was tabulated. The recombination frequency between different markers was used to narrow down the size of the target interval in which *rmr5* mutations were contained (Supplemental Table 4).

### Cloning of Target Loci and Identification of *rmr5* Lesions

Target gene models in the interval of interest were identified using the genome annotation tools available on Ensemble (http://ensembl.gramene.org/Zea_mays/Info/Index). Oligonucleotide primers for these models were generated using Sequencher (Gene Codes Corporation, Ann Arbor, MI) and Primer3 (http://bioinfo.ut.ee/primer3-0.4.0/primer3/). Genomic DNA from mutant individuals was amplified with these primers, and PCR amplicons were sequenced by the Vincent J. Coates DNA Sequencing Facility (UC Berekely). Comparison of amplified sequences to the reference sequence of the candidate gene models was performed with Sequencher and the online databases ChromDB (http://www.chromdb.org) and Ensemble (http://ensembl.gramene.org/Zea_mays/Info/Index).

Genomic DNA from the reference inbred A619 and individual homozygous *rmr5* mutants and candidate gene sequences were amplified using PCR. DNA samples from *rmr5-1*, *rmr5-2*, and *rmr5-3* homozygotes were extracted from embryos (Erhard et al., 2009), while A619 and *rmr5-4* samples were extracted from seedling leaves as described above. B73 cDNA was synthesized from high molecular weight RNA isolated from 4cm immature ears using Superscript III (Invitrogen) following the manufacturer’s protocol (Erhard et al., 2013). The thermocycling conditions for amplifying targeted regions were the following: 2 minutes at 94°C, 35 cycles of 30 seconds at 94°C, 30 seconds at 58°C, and 1 minute at 72°C, and a final extension of 10 minutes at 72°C. Primers used for sequencing are listed in Supplemental Tables 5 and 6. PCR products were cleaned with *Exonuclease*I (*Exo*I)(Fermentas; Pittsburgh, PA) and Shrimp Alkaline Phosphatase (SAP) (Promega; Madison, WI) incubated at 37°C for 30 minutes and 80°C for 15 minutes. Cleaned products were sequenced at the Vincent J. Coates DNA Sequencing Facility (UC Berkeley). Sequences were analyzed with Sequencher (Gene Codes Corporation, Ann Arbor, MI).

### Protein Analyses

The DCL3 conserved domains were identified using the Pfam (http://pfam.janelia.org; Punta et al., 2012) “Search Sequence” function with an E-value of 1.0. These same domains were also defined with other protein annotation databases: CCD (http://www.ncbi.nlm.nih.gov/Structure/cdd/wrpsb.cgi; Marchler-Bauer et al., 2011), PROSITE (http://prosite.expasy.org/; de Castro et al., 2006), SMART (http://smart.embl-heidelberg.de/; Letunic et al, 2012). Summary of the domains and their locations from the four databases are found in Supplemental Table 7.

For phylogenetic comparisons, DCL-type sequences from plant species were obtained through BLASTP searches on Phytozome (www.phytozome.net; Goodstein et al., 2012) using all *Arabidopsis thaliana* DCL protein sequences as queries. All sequences were queried through Pfam (http://pfam.janelia.org; Punta et al., 2012) to verify the annotation. Several sequences lacking certain DCL conserved domains were predicted with FGENESH+ (www.softberry.com) using similarity to AtDCL or OsDCL, unless otherwise noted. A full list of plants obtained and used in the phylogenetic analysis with the locus names for each plant DCL can be found in Supplemental Table 8. Monocot, *Arabidopsis thaliana, Aquilegia coerulea*, and *Volvox carteri* sequences were aligned with MAFFT (http://mafft.cbrc.jp/alignment/server/; Katoh and Toh, 2008) and a maximum likelihood tree was constructed with PHYML (http://mobyle.pasteur.fr/cgi-bin/portal.py; Néron et al., 2009) with the following conditions: 100 bootstraps sets to analyze, JTT amino acid substitution model, and NNI-SPR tree topology search operation. The tree was diagramed using DRAWGRAM via PHYLIP (http://evolution.gs.washington.edu/phylip.html; Felsenstein, 2005) and edited on Adobe Illustrator CS5.

### *In vitro* Transcription Reactions

Nuclei were isolated from 14-day post-imbibition seedling sheath sections (seedling region above soil and below first leaf blade) and used in *in vitro* transcription reactions with 32P-CTP as detailed in Erhard et al. (2009). Radiolabeled nascent RNAs were purified and hybridized to slot blots containing antisense riboprobes made to *pl1*, *a1*, *uq2*, and pBS (empty vector) as described in Hollick et al. (2005). Phosphorimaged results were processed using imageJ as described in Erhard et al. (2009) normalizing background-corrected values to *uq2* signals.

### DNA Methylation Assays

Bisulfite sequencing was carried out with gDNA isolated as in Dorweiler et al. (2000) from terminal flag leaves of heterozygous and homozygous *rmr5-2* mutant siblings. Five hundred nanograms of gDNA was bisulfite converted following manufacturer’s specifications using the EZ DNA Methylations kit (Zymo Research; Irvine, CA). Bisulfite-treated gDNA was amplified from −660 to +97 from the transcription start site of *Pl1-Rhoades* (Cone et al., 1993) using primers Doppia-bis-R primer: 5′ – GTGATTAGGTAGAAGTGGGAG – 3′ (Erhard et al., 2013) and 4R-pl1doppiabis: 5′ – CCCCTCTCTTCACCCCTTCCTT – 3′ (Gruntman et al., 2008). Gel purified (Qiagen, Qiaex II Gel Extraction Kit; Valencia, CA) ∼0.75 kb amplicons were cloned using the pGEM-T Easy vector system (Promega; Madison, WI) following manufacturer’s protocol and the plasmids were then transformed into *TOP10* cells (Invitrogen; Grand Island, NY). Individual plasmid clones were sequenced by the Plant-Microbe Genomics Facility (The Ohio State University) using the Doppia-bis-R primer (Erhard et al., 2013). Geneious (v6.1.8; Kearse et al., 2012) was used to trim sequences and validate sequence quality before bisulfite analysis was carried out with Kismeth (Gruntman et al., 2008). The parameters used for the Kismeth web suite were 0.8 min fraction of positive matches and 0.25 min fraction of length and the data included came from clones representing two *rmr5-2* heterozygotes and three *rmr5-2* homozygous mutants.

For the chop-PCR assays, terminal flag leaf gDNA samples used for the bisulfite analyses were assessed using methylation-sensitive restriction digests. Five microgram samples were individually digested overnight at 37°C with different methylation-sensitive restriction enzymes (*Hae*III, *Ava*II, *Hpa*II, *Sau*3AI and *Psp*GI) (New England Biolabs; Ipswitch, MA). A control digest was also prepared using *Mcr*BC for 2 hours at 37°C (Xie et al., 2012). The digested DNA was then PCR amplified using previously described primers (Lamb et al., 2007) for *CRM* (*gypsy*-like centromeric LTR) and *Prem-2* (*copia-*type LTR), newly designed primers (RGL_ubid-3F – AAGGAAATTAGGGGTGGGCG and RLG_ubid-3R – GCTGCAAGCGATGACATGAG) delimiting an upstream region of *ocl2* containing part of a *ubid gypsy*-type LTR, and the ANSSR-5 primer set (see Supplemental Table 3) specific for an arbitrary intergenic region of chromosome *1*. Undigested gDNA was amplified simultaneously as a loading control. The PCR conditions used were: 95°C for 2 m, 95°C for 1 m, 55°C for 1m, 72°C for 1m 30s, repeat for 30 cycles and 72°C for 1m.

### sRNA Libraries and Analyses

sRNA libraries were constructed, sequenced, and analyzed from total RNA isolated from 4.5cm developing cobs of *Rmr5* / *rmr5-2* and *rmr5-2* / *rmr5-2* siblings as previously specified (Barbour et al., 2012). Library statistics and initial analyses are presented in Supplemental Tables 9 and 10 respectively. Additional analyses of sRNAs mapping to a 100 Mb region of chromosome *1* are detailed in the Supplementary Methods and Supplemental Dataset 1. Briefly, sRNA islands consisting of 3 or more distinct reads overlapping by at least 1 base mapping to the first 100 Mb of chromosome *1* were computationally defined as clusters as described in Groszmann et al. (2011) and then analyzed for relative hits-normalized sRNA abundances.

### Primers Used in the Work

Oligonucleotide primer sequences are found in Supplemental Tables 3 (SSLP markers), 5 (*dcl3* genomic amplicons), and 6 (*dcl3* cDNA amplicons) or are described above.

### Accession Numbers

DNA sequence data from this work can be found in the GenBank/EMBL database under the following accession numbers: *dcl3-1* / *rmr5-1* (JX504671), *dcl3-2* / *rmr5-2* (JX504672), *dcl3-3* / *rmr5-3* (JX504673), *dcl3-4* / *rmr5-4* (JX504674). sRNA sequences have been deposited in the NCBI SRA database under accessions SRR16938685 and SRR16938686. The raw and normalized SBS data are also available at http://mpss.danforthcenter.org/maize.

## Supplemental Data

### Supplemental Methods

**Supplemental Figure 1.** Bulked Segregant Analysis for *rmr5-3*.

**Supplemental Figure 2.** Effects of ZmDCL3 on Cytosine Methylation at the *Pl1-Rhoades doppia* Boarder.

**Supplemental Figure 3.** Effects of ZmDCL3 at Repetitive Elements.

**Supplemental Figure 4.** Summer 2013 Morphometric Data.

**Supplemental Table 1.** Segregation Statistics for New ems-Derived Alleles.

**Supplemental Table 2.** Genetic Complementation Progeny Tests.

**Supplemental Table 3.** List of Oligonucleotide Primers Identifying SSLP Markers Used in Mapping Efforts.

**Supplemental Table 4.** Recombination Mapping Data.

**Supplemental Table 5.** List of Oligonucleotide Primers Used for Genomic DNA Amplifications and Sequencing.

**Supplemental Table 6.** List of Oligonucleotide Primers Used for cDNA Amplifications and Sequencing.

**Supplemental Table 7.** Locations of Domains Identified within ZmDCL3.

**Supplemental Table 8.** List of *dcl* Genes Among the Grasses and Outgroups.

**Supplemental Table 9.** Summary Statistics of SBS libraries.

**Supplemental Table 10.** Analysis of SBS Reads.

**Supplemental Dataset 1.** Chromosome *1* Cluster Analysis of SBS Reads.

## ACKNOWLEDGEMENTS

Thanks to the American Chemical Society Project SEED for supporting Glenna Kong who discovered three of the *dcl3* mutants. We are grateful to Drs. Blake Meyers and Stacey Simon who prepared and sequenced the small RNA libraries, Janelle Gabriel for both 5mC and morphometric analyses, and Emily McCormick for performing the run-on transcription experiments. NCBI, Gramene, and MaizeGDB provided critical sequence and data resources. Maize nurseries were supported in part by the University of California College of Natural Resources Oxford Facilities Unit, the Ohio Agricultural Research and Development Centers Waterman Agricultural and Natural Resources Laboratory, and The Ohio State University College of Arts & Sciences Biological Sciences Greenhouse. This work was supported by Syngenta, the National Research Initiative of the USDA Cooperative State Research, Education, and Extension Service (99-35301-7753, 2001-35301-10641, and 2005-35301-15891), the National Science Foundation (MCB-0419909, MCB-0920623) and The Ohio State Foundation. All *rmr* materials and their uses are covered by U.S. patent 8134047 assigned to The Regents of the University of California. The views expressed are solely those of the authors and are not endorsed by the sponsors of this work.

## AUTHOR CONTRIBUTIONS

J.B.H. designed the research. A.S.N., I.T.L., J.R.B.T., N.C.D. and J.B.H. performed research. A.S.N., I.T.L., J.R.B.T., N.C.D. and J.B.H. analyzed data. J.B.H. wrote the article.

