## Supplemental Materials for "A dicer-like3 protein affects paramutation at multiple loci in *Zea mays*"

### Supplemental Data

### Supplemental Methods

#### Genetic tests of new mutant alleles

Genetic test results (Supplemental Tables 1 and 2) show ems051109, ems051113, ems051271 and ems062989 mutations define recessive alleles of a novel locus designated *rmr5*. All four ems-derived mutations appeared to segregate as single locus recessive types in genetic tests (Supplemental Table 1). One homozygous ems051109 / ems051109 plant having a *Pl-Rh*-like phenotype (ACS 7) was outcrossed to an A632 *Pl′* / *Pl′* tester. All $F_1$ progeny plants (17 individuals) had a *Pl′*-like anther phenotype indicating the ems051109 allele is recessive and further that *Pl′* alleles transmitted from homozygous ems051109 / ems051109 plants are not rendered recalcitrant to paramutation. The *Pl-Rh*-like anther phenotype was recovered in 2/14 $M_2$ plants (Supplemental Table 1). The observed frequency of *Pl-Rh*-like phenotypes (14%) is not significantly different from the 25% expected from a single locus recessive mutation (Pearson's goodness of fit $X^2 = .64$; $p = .42$). One homozygous ems051113 / ems051113 plant having a *Pl-Rh*-like phenotype was outcrossed to three distinct *Pl′* / *Pl′* testers (A619, A632, W23). All $F_1$ progeny plants (33 individuals from 3 independent outcrosses) had a *Pl′*-like anther phenotype (13 ACS 1; 19 ACS 2; 1 ACS 3) indicating that the ems051113 allele is recessive and further that *Pl′* alleles transmitted from homozygous ems051113 / ems051113 plants are not recalcitrant to subsequent paramutation. The *Pl-Rh*-like anther phenotype was recovered in 2/23 $M_2$ and 71/348 $F_2$ plants (Supplemental Table 1). The observed frequencies of *Pl-Rh*-like phenotypes (9% and 20% respectively) are not significantly different from the 25% expected from a single locus recessive mutation (Pearson's goodness of fit $X^2 = 2.44$; $p = .12$; $X^2 = 1.82$; $p = .18$). One homozygous ems051271 / ems051271 plant having a *Pl-Rh*-like phenotype was also outcrossed to three distinct *Pl′* / *Pl′* testers (A619, A632, W23). All but three of $F_1$ progeny plants (29 individuals from 3 independent outcrosses) had a *Pl′*-like anther phenotype (17 ACS 1; 9 ACS 2). The remaining three had a complete tasselseed-like phenotype in which no anthers were produced. These data indicate that the ems051271 allele is recessive and further that *Pl′* alleles transmitted from homozygous ems051271 / ems051271 plants are not recalcitrant to subsequent paramutation. The *Pl-Rh*-like anther phenotype was recovered in 2/12 $F_2$ plants (Supplemental Table 1). The observed frequency of *Pl-Rh*-like phenotypes (17%) is not significantly different from the 25% expected from a single locus recessive mutation (Pearson's goodness of fit $X^2 = .5$; $p = .48$). One homozygous ems062989 / ems062989 plant having a *Pl-Rh*-like phenotype was outcrossed to a *Pl′* / *Pl′* plant (A632). All $F_1$ progeny plants (29 individuals) had a *Pl′*-like anther phenotype (17 ACS 1; 9 ACS 2). These data indicate that the ems062989 allele is recessive and further that *Pl′* alleles transmitted from homozygous ems062989 /ems062989 plants are not recalcitrant to subsequent paramutation. The *Pl-Rh*-like anther phenotype was recovered in only 2/37 (5%) $M_2$

plants (Supplemental Table 1) which was significantly different from the 25% frequency expected from a single locus recessive mutation (Pearson's goodness of fit $X^2$ = 5.68; *p* = .017). However, *Pl-Rh*-like anther phenotypes were recovered in 6/36 (20%) and 6/34 (18%) $F_2$ plants (Supplemental Table 1) which was not significantly different from the 25% frequency expected from a single locus recessive mutation (Pearson's goodness of fit $X^2$ = 1; *p* = .32 and $X^2$ = .74; *p* = .39, respectively). These segregation ratios are consistent with the hypothesis that the dark-anther phenotypes found in $M_2$ and $F_2$ progenies are due to single locus recessive mutations.

Results of complementation test crosses (Supplemental Table 2) indicate the ems051109, ems051113, ems051271 and ems062989 define loci distinct from *mop1*, *rmr1*, *rmr2*, *rmr6*, and *rmr7*. However, all four mutations fail to complement each other indicating that they define the same genetic locus. If one mutation is allelic to another, then 1/2 of all progeny from a given complementation cross between a presumed heterozygote and a homozygous parent should have a *Pl-Rh*-like anther phenotype. The observed frequencies of *Pl-Rh*-like types from these crosses are not significantly different from the expected frequency for this hypothesis (+/ems051109 X ems051113/ems051113, 53%, Pearson's goodness of fit $X^2$ = .0003, *p* = .97; +/ems051109 X ems062989/ems062989, 32%, Pearson's goodness of fit $X^2$ = 2.74, *p* = .098; +/ems051113 X ems062989/ems062989, 50%, Pearson's goodness of fit $X^2$ = 0, *p* = 1; +/ems051271 X ems062989/ems062989, 52%, Pearson's goodness of fit $X^2$ = .048; *p* = .83). All progeny plants generated by crossing parents homozygous for ems051113 and ems051271 also had *Pl-Rh*-like anthers as expected. The only pairwise combination not tested was between ems051109 and ems051271. These data indicate that ems051109, ems051113, ems051271, and ems062989 are alleles collectively defining a novel *rmr* locus, hereafter referred to as *rmr5*, that are renamed *rmr5-1*, *rmr5-2*, *rmr5-3*, and *rmr5-4*, respectively.

**Additional sRNA Analyses**

Reads aligning with no mismatches to the first 100Mb of Chromosome *1* using Bowtie (Langmead et al., 2009) (Bowtie [-f -v 0 -a –B 1]) were then aligned to the entire ZmB73 RefGen_v2 genome (Bowtie [-t -f -v 0 -a]) to calculate the total mappings for each read. These two bowtie outputs were merged into a working BED-formatted file and sorted (sort -k1,1 -k2,2n). Islands of 24-nt sRNAs were called from the ZmB73-uniquely-mapping *Rmr5* / *rmr5-2* het reads by combining 3 or more overlapping (>=1 bp) (Groszmann et al., 2011) using ClusterBed (BEDTools) (Quinlan et al., 2010) A total of 9,758 24-nt sRNA islands were identified in the first 100Mb of Chromosome *1*. sRNA islands were combined into clusters by adding a 100-nt pad to each island end and merging any now overlapping islands with mergeBed -d 100 (BEDTools). A total of 6,099 24-nt clusters were identified for which both read abundances and genome-wide hits-normalized abundances of 20, 21, 22, 23, 24, and 25-nt species were recorded for each cluster (Supplemental Dataset 1). Hits-normalized abundances per cluster nucleotide were then calculated and compared in Supplemental Dataset 1.

### Supplemental Figures

### Supplemental Figure 1

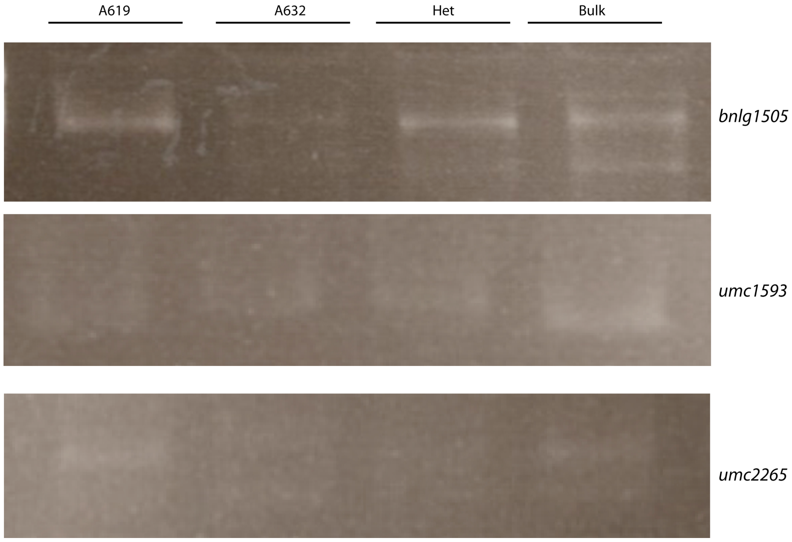

**Supplemental Figure 1.** Bulked Segregant Analysis for *rmr5-3*. Amplicons generated with the indicated SSLP markers were separated by agarose gel electrophoresis and stained with ethidium bromide. A619 and A632 represent genomic DNA from the respective parental lines. Het represents equal concentrations of A619 and A632 genomic DNA and Bulk represents genomic DNA pooled from 15 individual *rmr5-3* mutants.

**Supplemental Figure 2**

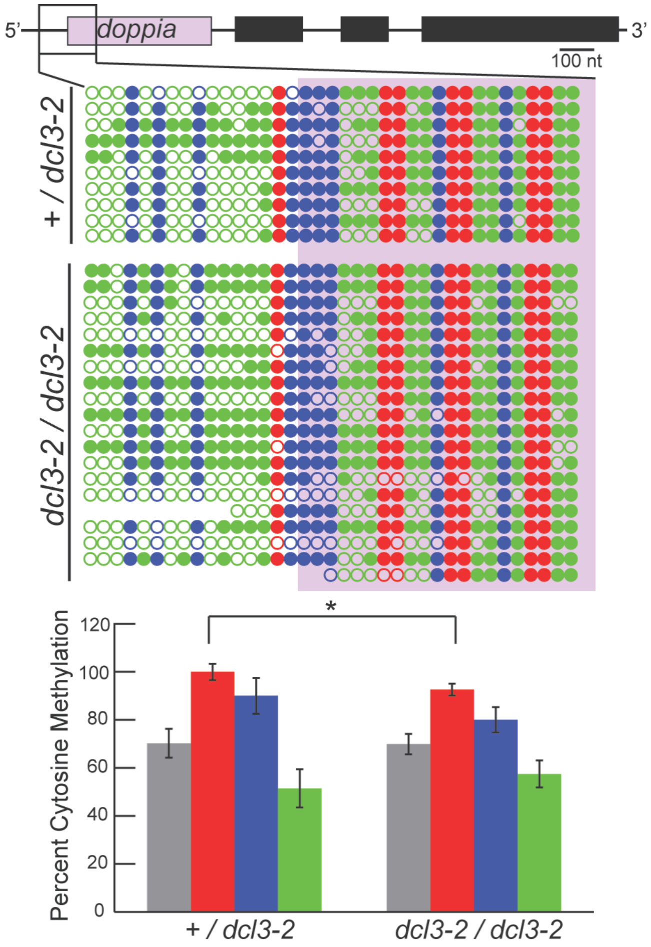

**Supplemental Figure 2.** Effects of ZmDCL3 on Cytosine Methylation at the *Pl1-Rh doppia* Boarder. A schematic for *Pl1-Rh*, with *doppia* sequences highlighted purple, and dot plots showing the methylation status at each cytosine within a region previously analyzed for the loss of RPD1 (Erhard et al., 2013). Dots and bars of red, blue, and green correspond with CG, CHG, and CHH cytosine methylation contexts. Grey bars correspond to total methylation. Mean methylation (± s.e.m.) was compared across the region for each methylation context between heterozygotes and mutants. Means were compared using a two-tailed Student's *t*-test; * - $P < 0.05$.

**Supplemental Figure 3**

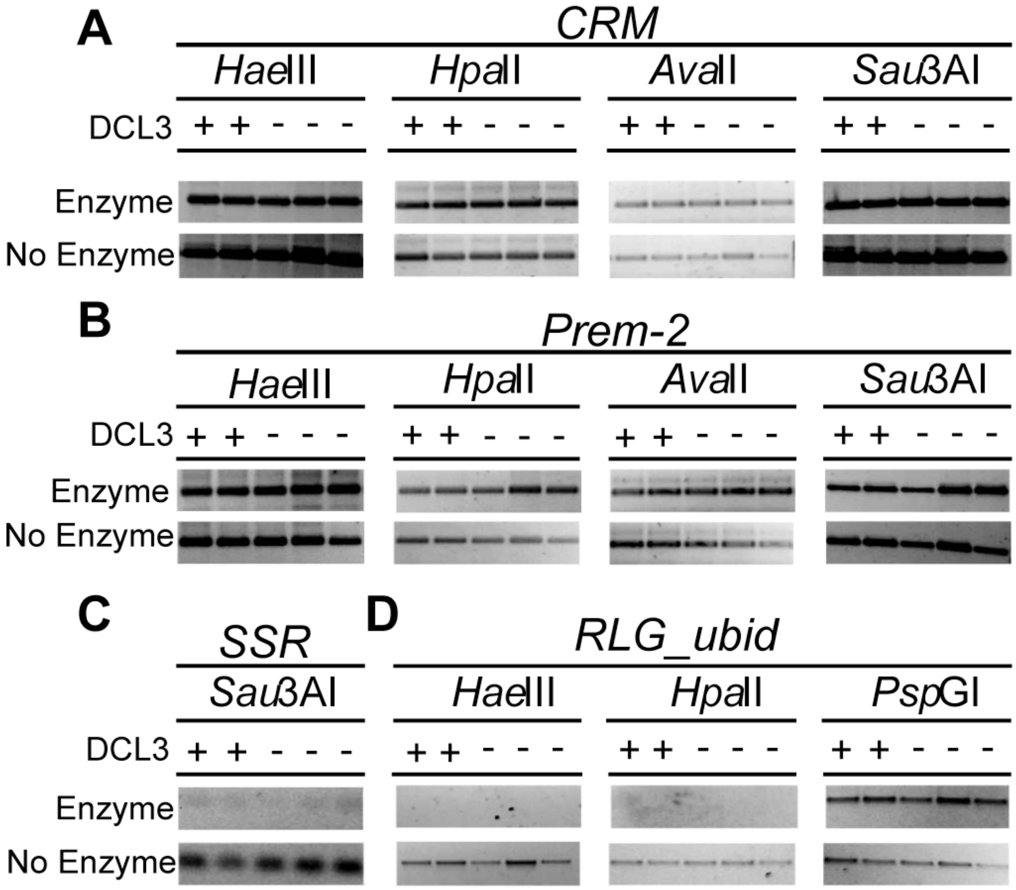

**Supplemental Figure 3.** Effects of ZmDCL3 at Repetitive Elements. Amplicons representing centromere-specific transposable elements (*CRM*; A), *copia*- like LTR retrotransposons (*Prem-2*; B), a satellite repeat region on chromosome *1* (*SSR*; C), and a *gypsy*-like LTR retrotransposon upstream of *ocl2* (*RLG_ubid*; D) were generated from genomic DNA samples prior to (No Enzyme) or following digestion with methylation-sensitive enzymes. *Hae*III and *Psp*GI digest CHH and CHG cytosine contexts*; Hpa*II and *Ava*II digest CHG and CG contexts; and *Sau*3AI digests all three cytosine methylation contexts. Sibling plants with (+) or (-) indicates presence or absence of ZmDCL3 function.

**Supplemental Figure 4**

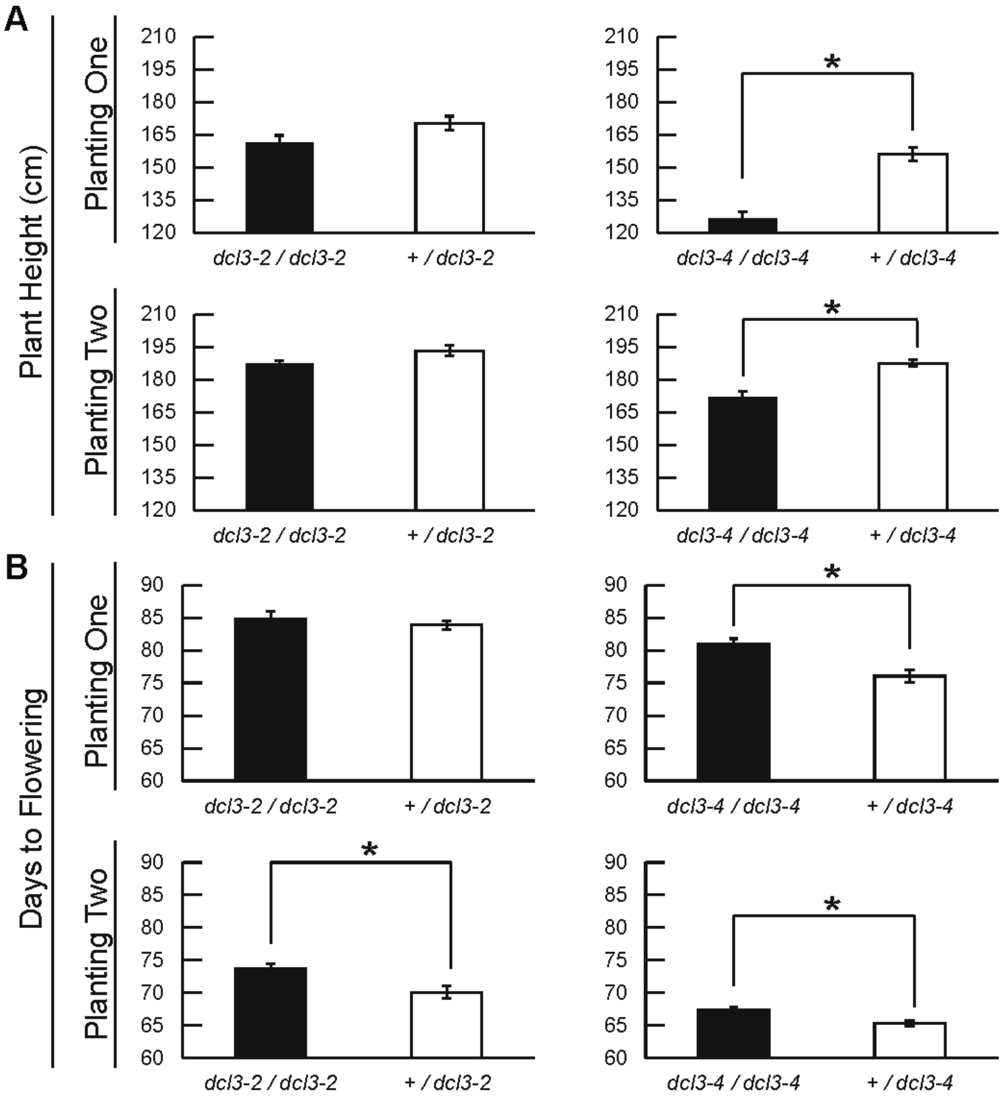

**Supplemental Figure 4.** Summer 2013 Morphometric Data. Progeny segregating 1:1 for either *dcl3-2* or *dcl3-4* mutants and respective heterozygotes were evaluated for plant height (A) and days to flowering (B). Two plantings for each progeny set were evaluated during the 2013 field season. Means for each trait (± s.e.m.) for the contrasting genotypes were compared using a two-tailed Student's *t*-test; *- $P < 0.05$. Planting 1: + / *dcl3-2* (n = 22); *dcl3-2* / *dcl3-2* (n = 16); + / *dcl3-4* (n = 20); *dcl3-4* / *dcl3-4* (n = 8). Planting 2: + / *dcl3-2* (n = 11); *dcl3-2* / *dcl3-2* (n = 10); + / *dcl3-4* (n = 31); *dcl3-4* / *dcl3-4* (n = 16).

### Supplemental Tables

#### Supplemental Table 1.
#### Segregation Statistics for New ems-Derived Alleles

| EMS allele | Progeny | | | | | | Statistics | |
|---|---|---|---|---|---|---|---|---|
| | Generation | Progeny Identifier | Number with specific anther color scores (ACS) | | | | Frequency of *Pl-Rh* types | *P* value[1] |
| | | | 7 | 5-6 | 1-4 | Total | | |
| 051109 | M2 | 034873 | 2 | 0 | 12 | 14 | 0.14 | n.s. |
| 051113 | M2 | 034877 | 2 | 0 | 21 | 23 | 0.09 | n.s. |
| 051113 | F2 | 072713 | 71 | 0 | 277 | 348 | 0.20 | n.s. |
| 051271 | M2 | 043043 | 2 | 0 | 10 | 12 | 0.17 | n.s. |
| 062989 | M2 | 062989 | 1 | 1 | 35 | 37 | 0.05 | * |
| 062989 | F2 | 072688 | 6 | 0 | 30 | 36 | 0.20 | n.s. |
| 062989 | F2 | 072691 | 3 | 3 | 28 | 34 | 0.18 | n.s. |
| Totals | | | 87 | 4 | 413 | 504 | 0.17 | n.s. |

[1] Pearson's goodness of fit analysis was applied to the null hypothesis that the observed frequency of progeny having a *Pl-Rh* / *Pl-Rh*-like phenotype (ACS 5-7) was unlike that expected from segregation of a recessive mutation; n.s. - not significant; * - significant ($P<0.05$).

**Supplemental Table 2.**
**Genetic Complementation Progeny Tests**

| EMS allele | Female Tester | Progeny | | | | | Statistics | |
|---|---|---|---|---|---|---|---|---|
| | | ID | Number with specific anther color scores (ACS) | | | | Frequency of *Pl-Rh* types | *P* value[1] |
| | | | 7 | 5-6 | 1-4 | Total | | |
| **051113** | **+/051109** | **070703** | **8** | **0** | **7** | **15** | **0.53** | **n.s.** |
| **051271** | **051113** | **070530** | **10** | **3** | **0** | **13** | **1** | **n.s.** |
| **062989** | **+/051109** | **070121** | **13** | **0** | **28** | **41** | **0.32** | **n.s.** |
| **062989** | **+/051113** | **070123** | **18** | **0** | **18** | **36** | **0.5** | **n.s.** |
| **062989** | **+/051271** | **070125** | **22** | **0** | **20** | **42** | **0.52** | **n.s.** |
| 062989 | *+/mop1-4* | 070108 | 0 | 0 | 13 | 13 | 0 | ** |
| 062989 | *+/rmr1-1* | 070109 | 0 | 0 | 8 | 8 | 0 | ** |
| *rmr1-5* | +/051109 | 070241 | 0 | 0 | 16 | 16 | 0 | ** |
| *rmr1-5* | +/051113 | 070154 | 0 | 0 | 13 | 13 | 0 | ** |
| *rmr1-5* | +/051271 | 070155 | 0 | 0 | 17 | 17 | 0 | ** |
| 062989 | *+/rmr2-1* | 070111 | 0 | 0 | 11 | 11 | 0 | ** |
| 062989 | *+/rmr6-1* | 070113 | 0 | 0 | 9 | 9 | 0 | ** |
| *rmr6-16* | +/051109 | 070306 | 0 | 0 | 29 | 29 | 0 | ** |
| *rmr6-16* | +/051113 | 070210 | 0 | 0 | 16 | 16 | 0 | ** |
| *rmr6-16* | +/051271 | 070211 | 0 | 0 | 9 | 9 | 0 | ** |
| 062989 | *+/rmr7-1* | 070183 | 0 | 0 | 14 | 14 | 0 | ** |
| *rmr7-3* | +/051109 | 070169 | 0 | 0 | 11 | 11 | 0 | ** |
| *rmr7-3* | +/051113 | 070096 | 0 | 0 | 27 | 27 | 0 | ** |
| *rmr7-3* | +/051271 | 070171 | 0 | 0 | 33 | 33 | 0 | ** |

[1] Chi-squared analysis was applied to the null hypothesis that the observed frequency of progeny having a *Pl-Rh* / *Pl-Rh*-like phenotype (ACS 5-7) was unlike that expected from genetic non-complementation; n.s. - not significant; ** - highly significant (*P*<<0.001).

**Supplemental Table 3.**
**List of Oligonucleotide Primers Identifying SSLP Markers Used in Mapping Efforts**

| name | sequence | Location (Mbp) |
|---|---|---|
| **Chromosome 1** | | |
| umc1354F | GATCAGCCCGTTCAGCAAGTT | 1.8 |
| umc1354R | GAGTGGAGGCGGAGGATCTG | 1.8 |
| umc1041F | CATTTCTTAGCACAACGCTGGTAAC | 6.1 |
| umc1041R | GCCACTGTGATTTCCCTTGTGT | 6.1 |
| bnlg1953F | CCTCGGAGCTCGATTTACAC | 23 |
| bnlg1953R | AACATTTAACCGCCGTCATC | 23 |
| bnlg1803F | GTATGCGTCGCTAGTCGTGA | 28.5 |
| bnlg1803R | TGTTGTCTATTGGCAACCGA | 28.5 |
| ANSSR-1F | ggggattccacttgtcattg | 36 |
| ANSSR-1R | tgtggcataaatatcctggtga | 36 |
| bnlg182F | AGACCATATTCCAGGCTTTACAG | 52 |
| bnlg182R | ACAACTAGCAGCAGCACAAGG | 52 |
| bnlg2180F | ACAAGGGCGTACCAACCAC | 53.5 |
| bnlg2180R | TGACCAGAGGCTTCCATACC | 53.5 |
| ASN21F | gaagcaagtgcagggtcaat | 56 |
| ASN21R | ctctcaggccgaggatactg | 56 |
| ASN23F | ttgcaaagtttaactgatggattt | 67 |
| ASN23R | cggaaacacaggtgatgagg | 67 |
| ASN24F | agcatcatcgtatcgacgttt | 75 |
| ASN24R | caaggagggtccccttacat | 75 |
| ASN25F | agatgcgcactgaactcctt | 80 |
| ASN25R | tcgcgtacgtggatgaaata | 80 |
| ASN27F | gaatcgtcacaaaaaggtcca | 90 |
| ASN27R | agccgtaggtttgagaacca | 90 |
| ASN29F | caactggaatgaaagcgttg | 110 |
| ASN29R | atgatatgctgtggcccttc | 110 |
| bnlg1057F | TTCACCGCCTCACATGAC | 128 |
| bnlg1057R | GCAACGCTAGCTAGCTTTG | 128 |
| umc1395F | TGAATGAGTGGCATTCAAAATCTG | 164 |
| umc1395R | CAGATTGCATGTGTGAGTGTGTGT | 164 |
| bnlg100F | TGCACGCACGGGCACTGAAC | 184.76 |
| bnlg100R | TAAGACATCTATGGCCACCGGAG | 184.76 |
| bnlg1273F | AAACACCAAACGTCACGTGG | 194.55 |
| bnlg1273R | GGCGACGAGATACAGGATGT | 194.55 |
| umc1035F | CTGGCATGATCACGCTATGTATG | 195.05 |

| | | |
|---|---|---|
| umc1035R | TAACATCAGCAGGTTTGCTCATTC | 195.05 |
| ANSSR-2F | cctgcaaccacagcaatactt | 217 |
| ANSSR-2R | gcctgtgttgacgcataatc | 217 |
| ANSSR-3F | tgcctacaagggagggtaat | 226 |
| ANSSR-3R | gagcctccgctcttttctct | 226 |
| ANSSR-5F | caggagctccagttcctcac | 236 |
| ANSSR-5R | ggtcatgcagtctcgcttct | 236 |
| ANSSR-8F | | 252 |
| ANSSR-8R | gtgggactcacgaagggata | 252 |
| umc2149R | AGCAGCACCATCGTAATAAGCAC | 272.11 |
| umc2149F | TACATGCAAAGCTAGCTAGTCGGA | 272.11 |
| bnlg667F | CGTGGATGTAAGGGGGCGCGCT | 290 |
| bnlg667R | GGCCGCTGCTCAACACAGGCAG | 290 |
| **Chromosome 2** | | |
| bnlg2042F | TGTCGCGTACTCGCATTTAG | 9.4 |
| bnlg2042R | TTTGATTGGTGATCTCGCAG | 9.4 |
| umc1261F | AGAAGTGCGTATGCTACAGTGGTG | 13.5 |
| umc1261R | CCTAGTGGTGGAGTTCTAGGCAAA | 13.5 |
| umc1845F | TGGTTGAACTGTTAAATCTGTCCTGA | 19.064 |
| umc1845R | TGGTAACCAGATTCCCACAGATG | 19.064 |
| ANSSR-9F | tcaggattgtgtctccatcg | 32 |
| ANSSR-9R | gccgctagcaagctgtatct | 32 |
| ANSSR-10F | gtcatcaacgcgctcaact | 36 |
| ANSSR-10R | cctgccacgtttttctctct | 36 |
| ASN34F | ctgcaccgacaaatgaagaa | 46 |
| ASN34R | ctgctcctcctggacttgag | 46 |
| ANSSR-13F | tcaatgttggttgatgtttgg | 62 |
| ANSSR-13R | ggaagaacatgagtcttctttgc | 62 |
| ASN41F | gtctccgtaatcggtcgttg | 80 |
| ASN41R | cggtctgcctttaggtgtgt | 80 |
| umc1635F | GCTGAGCAGATCTTTCCTTGTTTC | 82.5 |
| umc1635R | AAGGAGCAGAACTCGGAGACG | 82.5 |
| umc1749F | CTTCATTTTTGTTGTTTCCCTGCT | 101 |
| umc1749R | CGTAAAAGACTGGATTGCCAAAAG | 101 |
| umc1028F | CCCAGGTAAAATTCGCTAGCCT | 145 |
| umc1028R | GGAACAAGGAAAGCTGAATACACG | 145 |
| bnlg1831F | TCGCTCATTTGCATACACCT | 149 |
| bnlg1831R | TAGGAACATGCCAGCAGTTG | 149 |
| bnlg1396R | TGCTTGAGTCGTCGAATCTG | 166 |
| bnlg1396F | CGCATTTCTCCTGCAGTACA | 166 |
| umc1875F | AAGCGACAGTTGGTTTCAGTTTTC | 168 |

| | | |
|---|---|---|
| umc1875R | AGTTGCATGACCTTGTCAACCTTT | 168 |
| bnlg1887F | CGAACCACTGTAGGCATGTG | 172 |
| bnlg1887R | ATCATGCAGAGCAGATGCAG | 172 |
| bnlg1233F | GAACACCAGAGGAGAGTGGG | 206.5 |
| bnlg1233R | TTCACTTGTCCACCACTGGA | 206.5 |
| bnlg1316F | CGAAACAGAGCCCAAAAGAC | 209.4 |
| bnlg1316R | GATCCGCGTCTAGCCCCT | 209.4 |
| mmc0381F | GTGGCCCTGTTGATGAG | 214.3 |
| mmc0381R | CGACGAGTACCAGGCAT | 214.3 |
| umc2144F | CCAGCCCCTATCTATTTGCTTGT | Chr5? |
| umc2144R | GAATACTATATCACGGTCGGTCGG | No BLAST record |
| **Chromosome 3** | | |
| bnlg1144F | TACTCGTCGTGTGGCGTTAG | 5.22 |
| bnlg1144R | AGCCGAGGCTATCTAACGGT | 5.22 |
| bnlg1647F | CGTCGTCTGTGGACGTACTG | 8 |
| bnlg1647R | AGAAGCTCACAAGCCTGCTC | 8 |
| umc1458F | CCAATAAACAAATCATCTCCCCCT | 8.015 |
| umc1458R | TGCTATGCTATGTACAGGGACAGG | 8.015 |
| ASN42F | tgcagttgttcttccgagtg | 10 |
| ASN42R | gtggtcgatgctgtggtaaa | 10 |
| ASN46F | agcccagacaatgcagagat | 35 |
| ASN46R | ccaagcatgacagtcaccaa | 35 |
| ASN47F | gatgatgatgacaacgacga | 42 |
| ASN47R | actgatatgcacccccagac | 42 |
| ASN48F | ctcttatcgtggccttcgac | 50 |
| ASN48R | tcgattgttggcgtgactaa | 50 |
| bnlg1638F | CATATCTCTAGCTTCTCGTCTTCG | 57 |
| bnlg1638R | ACACCGATCGAGGAAGAATG | 57 |
| ANSSR-15F | tgaaaaatatccatatccgtatcc | 72 |
| ANSSR-15R | caagattaggggtgaaaacg | 72 |
| ANSSR-16F | tgggctttcaaatcaacaaa | 76 |
| ANSSR-16R | ttatatgaacccgccaaaaa | 76 |
| ANSSR-19F | gccactttgcagtagtgcttt | 92 |
| ANSSR-19R | cgcggttacagtatcagcaa | 92 |
| ANSSR-20F | ttcaatctgtatttccccatgtt | 96 |
| ANSSR-20R | attgccccttgtcagcttag | 96 |
| bnlg1628F | GTAGGGTTCAAGGAGGCACA | 115 |
| bnlg1628R | CTCTCTGGTGAGCTGGCTTT | 115 |
| bnlg1447F | GAGAGGAGAGGCTGAGCTGA | 122 |
| bnlg1447R | TCCTCCCACTGAATTTCCAC | 122 |
| mmc0022F | AGGTGTTGTTTTTGTTCGCT | 140 |

| | | |
|---|---|---|
| mmc0022R | TGCTTGTTTAAGCTCATTATT | 140 |
| ANSSR3-141F | ctccggcataaggtattcca | 141 |
| ANSSR3-141R | cgtgtaaggccttcgatctc | 141 |
| ANSSR3-142F | ctcaggaagggggatctgtt | 142 |
| ANSSR3-142R | cactgcttctcgatgtccaa | 142 |
| ANSSR-147-1F | agctgggatacaagcaggag | 147 |
| ANSSR-147-1R | gacaaggcgaagttggaaag | 147 |
| bnlg1505F | GAAAGACAAGGCGAAGTTGG | 147 |
| bnlg1505R | GCTTCTGAACTGGATCGGAG | 147 |
| ANSSR-149-1F | catgtacaggagccgactgg | 149 |
| ANSSR-149-1R | cctcttttgtggagccaaat | 149 |
| ANSSR3-150F | acatgctgcaggtcctctct | 150 |
| ANSSR3-150R | tcgcacgctacttccactta | 150 |
| ANSSR-151-1F | aagcccaaaattccttcgat | 151 |
| ANSSR-151-1R | gtccgacctagccttcctct | 151 |
| ANSSR3-154F | tgcgtgcacatgctttttac | 154 |
| ANSSR3-154R | cctgcctctttttgctatgc | 154 |
| ANSSR3-156F | cggcgagaaactgaactga | 156 |
| ANSSR3-156R | ccgcctgctcgtagtaaaac | 156 |
| ANSSR-157-2F | tcctggctaggacttccatt | 157 |
| ANSSR-157-2R | ttcccaatatgcttcttctgc | 157 |
| ANSSR3-158F | tgtcctctagcactggcattt | 158 |
| ANSSR3-158R | tacgttccgcagcaacaata | 158 |
| ANSSR-159-1F | cgtagtaatttgtatcgtgaaattagg | 159 |
| ANSSR-159-1R | ttcaacaacatcctttaaatttgg | 159 |
| umc2265F | CTGGACGTGGACTCAGACACC | 159 |
| umc2265R | AAGACGGTCCCGAAGAAAGC | 159 |
| ANSSR3-160F | cagcaaagcgctgtatgtgt | 160 |
| ANSSR3-160R | cttcgggacgaggaacgtat | 160 |
| ANSSR-161-2F | gaggaatttggtaggcagca | 161 |
| ANSSR-161-2R | ccatccattatgcccaatct | 161 |
| ANSSR3-163F | ggcatggacgtcatagtcct | 163 |
| ANSSR3-163R | ggtgacacggcttctctctc | 163 |
| ANSSR3-164F | tgccgagaggagagaaacat | 164 |
| ANSSR3-164R | tgtgcaacaaggagacaagc | 164 |
| ANSSR-165-1F | attaccacagacgacgacga | 165 |
| ANSSR-165-1R | cgaattttattttgcgacaga | 165 |
| ANSSR3-166F | tgtttgatgaggctctgacg | 166 |
| ANSSR3-166R | gtgccaatgtgaagcaagaa | 166 |
| umc1593F | CATGTTGATCATATGCACGAGAGA | 167 |
| umc1593R | CAGCCTGGTGACTCATGGTTAAT | 167 |

| | | |
|---|---|---|
| phi102228F | ATTCCGACGCAATCAACA | 178 |
| phi102228R | TTCATCTCCTCCAGGAGCCTT | 178 |
| ANSSR-21F | cgccaagaagaaacacatca | 190 |
| ANSSR-21R | cctgtgaagcctatggagga | 190 |
| ANSSR-23F | aagctatttggatgtctgcatct | 200 |
| ANSSR-23R | tgcactagagctaatactccttcc | 200 |
| ANSSR-24F | actccctgcaagcgtaaaaa | 205 |
| ANSSR-24R | atggcatcctgtattgtgct | 205 |
| ANSSR-25F | gtcgagttcgaagcccaat | 210 |
| ANSSR-25R | atagcggggaagcaggtagt | 210 |
| umc1639F | CTAGCCAGCCCCCATTCTTC | 228.9 |
| umc1639R | GCAAGGAGTAGGGAGGACGTG | 228.9 |
| **Chromosome 4** | | |
| bnlg1370F | TATTTAATTTAGTGTGGAGCTCACG | 1.547 |
| bnlg1370R | CGAGGGTCAGTTGTTGCTCT | 1.547 |
| bnlg1318F | TTATGTGTGCAGAACGACTCG | 4.915 |
| bnlg1318R | AGCATGGCAGAGAAGGTGAT | 4.915 |
| phi096F | TCCACCATTTGACACTTAGGCA | 29.9 |
| phi096R | GCGTAGGACGACCGTTGAA | 29.9 |
| nc005F | CCTCTACTCGCCAGTCGC | 36.7 |
| nc005R | TTTGGTCAGATTTGAGCACG | 36.7 |
| ANSSR26F | gaaggtttgacctggtgcat | 40 |
| ANSSR26R | aattattccgaaaacccgaaa | 40 |
| ANSSR27F | gtgccaaaccatgtctagga | 47 |
| ANSSR27R | ctcccgatatacctgcccta | 47 |
| ASN49F | aagaaggcggatctgacaaa | 53 |
| ASN49R | ctctctcgctgctcagtgg | 53 |
| ASN53F | ctcccaccgaacacaaatct | 70 |
| ASN53R | aaaaccatggaggacacgac | 70 |
| ASN53F | aaaaccatggaggacacgac | 70 |
| ASN53R | cctcagactgatgggcagat | 70 |
| umc1926F | ATGCCAGCATTCTTCATCCTACAT | 80.5 |
| umc1926R | TGAGGCTTGGTCCACTAAAGAAAG | 80.5 |
| ANSSR31F | ggctacttgcagtggctttt | 98 |
| ANSSR31R | acatccttgctccgtgaaac | 98 |
| ANSSR33F | gggcctacaagaagatggtg | 117 |
| ANSSR33R | cttgcgaaggaagggttgt | 117 |
| bnlg1755F | CCTAGTAGACCTCACCGCCA | 117.18 |
| bnlg1755R | GGAGTTCACCGATGGCAC | 117.18 |
| ASN56F | ggcataatgctgatggcata | 120 |
| ASN56R | accgcttcttcattcacagg | 120 |

| | | |
|---|---|---|
| ASN57F | aaagaagaacgatggtggaa | 125 |
| ASN57R | gagggagaggaaggcagtct | 125 |
| ASN58F | ggtgtggctatgctgttgaa | 135 |
| ASN58R | ttttgttttccgtgcatcct | 135 |
| ANSSR35F | gtcccgcaattcatttcct | 145 |
| ANSSR35R | tggcaccatggtattctcaa | 145 |
| ASN61F | tgcagaaataaaccgagttacg | 156 |
| ASN61R | ccaaacacatatgaccaaacaca | 156 |
| umc1043F | CTGCGCATGAGAAAACAGTGG | 180.9 |
| umc1043R | TCCATCTCTGGCTGGAGGTTAAT | 180.9 |
| ASN67F | aagggtggtcctggtgatg | 205 |
| ASN67R | cctggtcgaggtgaaggtt | 205 |
| ASN68F | tttgcaacagtgccgattac | 210 |
| ASN68R | gaatacatgccgcgattaca | 210 |
| umc1101F | GCTGAAAAACGGAGTTCATATGGT | 241.8 |
| umc1101R | AAGCTTATCCACCTCGAGGAAAAC | 241.8 |
| umc1503F | TTCATGACACACAAACCACAGATG | 242.5 |
| umc1503R | GCACCCTAGCAGACTACAACATCC | 242.5 |
| umc1058F | AGCAAGCAGTTCGAAACAAGGAT | 245.6 |
| umc1058R | GACACCAGCACCACTTGAACG | 245.6 |
| ASN69F | tccacctccatcaggaactc | |
| ASN69R | tccaaaattcccaaaacatca | |
| **Chromosome 5** | | |
| umc1416F | AGATGAATGTTGGGGTCAACAAGT | 10 |
| umc1416R | CTTGTCAGCCACAGAAGTGCC | 10 |
| umc1097F | CTCGTCAACGTCAACCCAAGTAAG | 18.7 |
| umc1097R | CTGTTAGATGTGCGACAACAGAGC | 18.7 |
| ANSSR37F | cgcactgaactcttcgttga | 22 |
| ANSSR37R | ccaccagatgacaggatgaa | 22 |
| ANSSR38F | ccgaaagcctgcttgagtc | 26 |
| ANSSR38R | ttaacgtgagacgaccgttg | 26 |
| ASN70F | cagcgaggcactaacctagc | 30 |
| ASN70R | gtggcttctcctccagcat | 30 |
| ANSSR41F | ggctccaaactctcaacagg | 42 |
| ANSSR41R | tgagaacagattcagatgtctatcg | 42 |
| ASN73F | tccttgtcatttatttatggggtaa | 46 |
| ASN73R | cctccaaggaagcacctgta | 46 |
| ASN74F | gaaagcccggagctcttaat | 50 |
| ASN74R | ttcgtttcgatcacaatcca | 50 |
| ASN75F | cggagtggagttccaggtaa | 57 |
| ASN75R | tcttctctgcctgtcgaggt | 57 |

| | | |
|---|---|---|
| ANSSR42F | gcacatcagttgcaagaagg | 66 |
| ANSSR42R | aggattttccctcattgatcc | 66 |
| ANSSR44F | accgaacctagtccagctca | 76 |
| ANSSR44R | cccatgccagtaccatctct | 76 |
| ANSSR45F | cgcaagaagttccccaaaa | 81 |
| ANSSR45R | tcgtccaccaataaagtctcg | 81 |
| ANSSR46F | gctccagttcatgcaaacaa | 86 |
| ANSSR46R | gacacgatcaacttgcaaca | 86 |
| umc2036F | TCAATCAAGCCTCTCGTAAGGAAC | 96 |
| umc2036R | CTCTTGATCTCAACCGAAATCCTG | 96 |
| ANSSR47F | ccaggcacaacctcaaaaat | 105 |
| ANSSR47R | ggtaatggtagggcggagtt | 105 |
| umc1557F | CTAGTTCTCTCAAAAGCGCCG | 105 |
| umc1557R | ACAATAGGTAGGTCCCCTGCTTT | 105 |
| ANSSR48F | ggtcccttccatagccataa | 111 |
| ANSSR48R | ccggaaaatcagctagtcca | 111 |
| ANSSR49F | tcgcaggatttggatctctt | 117 |
| ANSSR49R | gaggagcctagctcttgtgc | 117 |
| bnlg603F | CTGAGCTGGCCCCTGTGAATGGTG | 149.9 |
| bnlg603R | CGCCCTCCGCTGCGCTTCTCT | 149.9 |
| umc1221F | GCAACAGCAACTGGCAACAG | 168.07 |
| umc1221R | AAACAGGCACAAAGCATGGATAG | 168.07 |
| umc1524F | TACAAGTAAACACGCGCAGGAGTA | 196.32 |
| umc1524R | TGTTTGAGCGACTTACTTGACCTG | 196.32 |
| bnlg609F | GCTCGTTCTCGCCAGTGTGCCG | 198.956 |
| bnlg609R | GGCCCGAGCCATCTCTGCTGC | 198.956 |
| umc1792F | CATGGGACAGCAAGAGACACAG | 211.8 |
| umc1792R | ACCTTCATCACCTGCAACTACGAC | 211.8 |
| umc1225F | CTAGCTCCGTGTGAGTGAGTGAGT | 212.4 |
| umc1225R | TTCCTTCTTTCTTTCCTGTGCAAC | 212.4 |
| umc1829F | GTTGATTGGTTGATGTGGAAACAA | 214.2 |
| umc1829R | CAGTTTGATGTTCATGGCTCTCTC | 214.2 |
| **Chromosome 6** | | |
| bnlg1600F | | 1.93 |
| bnlg1600R | TAGGCATGCATTGTCCATTG | 1.93 |
| bnlg1432F | AAAGCAAACAAACAATGGGC | 19.2 |
| bnlg1432R | TGCGTGCAGTGACATATTCA | 19.2 |
| bnlg1867 | CCACCACCATCGTAGGAGTT | 19.5 |
| bnlg1867 | CAGTACACAGCAGGCAGCTC | 19.5 |
| bnlg249F | CCGGTCGCAGTTAGTAGATGAT | 58.6 |
| bnlg249R | TCGGCGTTGATTTCGTCAGTA | 58.6 |

| | | |
|---|---|---|
| bnlg1422F | GACGATTAACAGGTGGGGAC | 83 |
| bnlg1422R | ATGATGCAAATGAGGCACAA | 83 |
| umc1887F | CTTGCCATTTTAATTTGGACGTTT | 102.9 |
| umc1887R | CGAAGTTGCCCAAATAGCTACAGT | 102.9 |
| bnlg1732F | AACTTTTGGCATTGCACTGG | 151.9 |
| bnlg1732R | CGTAAGTGCACACGGCATTA | 151.9 |
| dupssr15F | GAAGTCGATCCATCCACC | 162.6 |
| dupssr15R | GGGGTAGTGGAGATAACTAGTG | 162.6 |
| umc1248F | CTTTGTCCATCGGCTTTATTCTTT | 164.6 |
| umc1248R | CACATTAAGTTACAAATACAAATCACCG | 164.6 |
| bnlg1740F | TTTTCTCCTTGAGTTCGTTCG | 164.7 |
| bnlg1740R | ACAGGCAGAGCTCTCACACA | 164.7 |
| umc1127F | GGTCCAGTGACATCTCAAAATGAA | 168.8 |
| umc1127R | ATATTCCCCCTCCCTAATTTTGCT | 168.8 |
| **Chromosome 7** | | |
| umc1036F | CTGCTGCTCAAGGAGATGGAGA | 10.1 |
| umc1036R | GACACACATGCACGAGCAGACT | 10.1 |
| mmc0162F | CGAAATCAAATCAAAACCATA | 19.2 |
| mmc0162R | CCTCCTATCGCTAGCTCGATC | 19.2 |
| umc1339F | GCACGGTTTTGTTGAATATGTGTG | 29.8 |
| umc1339R | GGTGGGAAGATATGCAACTGTCTC | 29.8 |
| umc1409F | GCTAGTAGACATCGACGGATCGAC | 46.35 |
| umc1409R | ATGACGTCCAGGAGGATGACC | 46.35 |
| umc1666F | TTATTGCCCTCCCTGTTCTTGT | 50 |
| umc1666R | ACCTTGACGCAGCAATCCTC | 50 |
| bnlg1380F | ACAATTCGATCGAGAGCGAG | 72 |
| bnlg1380R | CCTTTCTTGCTGGTTCTTGC | 72 |
| umc1585F | CGGCCTATGTAACAATCCCTAGC | 115 |
| umc1585R | AAGGGAAAGAATAATCCAACCGTC | 115 |
| umc1001F | GCTACCCGCGGACATATAAT | 141.9 |
| umc1001R | CCATGGGTAAAAACCCTACAGTG | 141.9 |
| umc1134F | AAAACTAACAGGCAGCAGACCAAC | 142 |
| umc1134R | ATCAGCAAGTGACTGAATTCCTCC | 142 |
| umc1154F | CCACCACAAGACAAGACAAGAATG | 167.5 |
| umc1154R | CCTGATCGATCTCATCGTCGT | 167.5 |
| umc1407F | AGGCTTACCTCCTGAGAAGCAGTT | 168.1 |
| umc1407R | AGGCTTAGCATCGGTGGAGAG | 168.1 |
| **Chromosome 8** | | |
| ASN79F | taggtcgctgaaggtcgttt | 1 |
| ASN79R | caagatcatgcatgctaggc | 1 |
| ASN81R | ttctcctccacgcatacaca | 10 |

| | | |
|---|---|---|
| ASN81F | ttcgggccctaatacaacag | 10 |
| ASN83F | ccttatcttccactacagagcgta | 20 |
| ASN83R | catgttgattgtttgcacca | 20 |
| bnlg669F | GCACGCACCAGCAGTCGGCAGT | 22 |
| bnlg669R | CGGCCTAGTGGGCATGGAGCCT | 22 |
| ASN84F | ggaaacggtacacgctaagg | 30 |
| ASN84R | cctgttttgtttcccactcaa | 30 |
| bnlg2082F | GACGGAAGGTGGAGCATAGA | 36 |
| bnlg2082R | ACGAACGTGATACGGGTCTC | 36 |
| umc1157F | AACTCGCTATCGAAAAACCACAAG | 69 |
| umc1157R | TCGGATTTTAGCTGAGCTTGTACC | 69 |
| umc1617F | GATGCACCACAGTAGAGGAGGAAT | 77 |
| umc1617R | GAAGATGAGGTTCAGGAAGGACAA | 77 |
| umc1302F | ATTTATTCAAACCGACGAAGCAA | 87 |
| umc1302R | TACAAGCGCTACTGCGATGTCTT | 87 |
| bnlg1863F | GGCGTTCGTTTTGCACTAAT | 90 |
| bnlg1863R | CGACACAGTTGACATCAGGG | 90 |
| phi115F | GCTCCGTGTTTCGCCTGAA | 99 |
| phi115R | ACCATCACCTGAATCCATCACA | 99 |
| umc1858F | GTTGTTCTCCTTGCTGACCAGTTT | 111.2 |
| umc1858R | | 111.2 |
| umc1005F | TTTGATCACAGACTTATCCCTGTT | 169.1 |
| umc1005R | CTAATGACGAACCCCTAAAAGGT | 169.1 |
| phi015F | GCAACGTACCGTACCTTTCCGA | 171.8 |
| phi015R | ACGCTGCATTCAATTACCGGGAAG | 171.8 |
| **Chromosome 9** | | |
| umc1279F | GATGAGCTTGACGACGCCTG | 1.59 |
| umc1279R | CAATCCAATCCGTTGCAGGTC | 1.59 |
| bnlg1583F | ATCAAGCTTATCGAGAGAGAGAGAG | 8.44 |
| bnlg1583R | CGACGGTGGAAAGACTGC | 8.44 |
| bnlg1730F | GGGTGCTCGTAGTAGGGGTT | 11.65 |
| bnlg1730R | AACACGTCAACAAGGGGAAG | 11.65 |
| umc1183F | ATGTCATTTTTGGCTTCTCGAAAT | 16.7 |
| umc1183R | GCATGTACACAACACAACCTTTCA | 16.7 |
| umc1586F | TAGGAGATGAGCTCGTCGGATAAG | 25.2 |
| umc1586R | GAGGATGAGGAGGATGGTAATGGT | 25.2 |
| ASN84F | ggaaacggtacacgctaagg | 30 |
| ASN84R | cctgttttgtttcccactcaa | 30 |
| ASN88F | gcattgcgggttttctttta | 51 |
| ASN88R | agcccgactgtctctagcaa | 51 |
| ASN89F | agacccagtccgaggtcttt | 62 |

| | | |
|---|---|---|
| ASN89R | ttggaggttatttgtttgtgc | 62 |
| ASN90F | gtgttcgcaatgtcctcctt | 72 |
| ASN90R | gcggtagaagcacacatgaa | 72 |
| umc1267F | TTACAACACGCATGCATCTAGCTT | 99.39 |
| umc1267R | AACAACAAAGAACTCACCAGCCTC | 99.39 |
| umc1120F | GCACAGCTTTGTCCCTCGAA | 117.3 |
| umc1120R | GACCCTCGAGCTTCTCTCATCTTT | 117.3 |
| umc1078F | AGGCACTAGCAGGCGAGAGG | 124.12 |
| umc1078R | GCGTAGTAACATCCATCCAACCAA | 124.12 |
| bnlg1270F | TAGTTAACATGAGCAAATTAACAAGA | 125.15 |
| bnlg1270R | TAGAAATGCAGAACCAGGGC | 125.15 |
| umc1231F | CTGTAGGGCTGAGAAAAGAGAGGG | 128.87 |
| umc1231R | CGACAACTTAGGAGAACCATGGAG | 128.87 |
| asn7F | cccagtcagtgtcggtgtaa | 134.2 |
| asn7R | gagcctaacacttgcgtatgg | 134.2 |
| asn8F | atgcttccatagccagcaac | 135.8 |
| asn8R | cggtttcagaactgcgtatg | 135.8 |
| umc1733F | ACACCCAACCTCCCACTGTAAAA | 140.91 |
| umc1733R | GATTGGGATTGGGATTGGAAAT | 140.91 |
| umc1366F | GTCACTCGTCCGCATCGTCT | 141.8 |
| umc1366R | CCTAACTCTGCAAAGACTGCATGA | 141.8 |
| asn15F | cgcccatctaaaccctaaca | 142.7 |
| asn15R | atcttggtgacatgctcgtg | 142.7 |
| umc1789F | ACCTCTCCTTTTTCCTCGCCTT | 143.45 |
| umc1789R | GTCAGAGAAGAGGCCGGGTC | 143.45 |
| umc1137F | ATCAGTCACTCTTCTGCCTCCACT | 148.97 |
| umc1137R | GGCTGGATAATGTTGTAGCTGGTC | 148.97 |
| umc1277F | TTTGAGAACGGAAGCAAGTACTCC | 150.45 |
| umc1277R | ACCAACCAACCACTCCCTTTTTAG | 150.45 |
| asn21F | actcctcggatgaggaggac | |
| asn21R | gaagaccagtggcacctagc | |
| **Chromosome 10** | | |
| phi041F | TTGGCTCCCAGCGCCGCAAA | 2.1 |
| phi041R | GATCCAGAGCGATTTGACGGCA | 2.1 |
| umc1337F | TGGATCTTTTATTTATGTTTTTATTCGG | 8.155 |
| umc1337R | CTGCCTGTAACGAATATGAATGC | 8.155 |
| ASN96F | cgtgatgttgagcctatgga | 37 |
| ASN96R | ccgatcgtgaaggaaggtaa | 37 |
| ASN93F | gacgtcgactttttcgaagc | 37 |
| ASN93R | ggcttctacgaagcaccatt | 37 |
| umc1179F | AGCTCCCCATCTTAATCCGTAAA | 75.5 |

| | | |
|---|---|---|
| umc1179R | ATTGAGCTCGGCGTAGAAAAA | 75.5 |
| umc1336F | GTACAAATGATAAGCAAGGGGCAG | 86.32 |
| umc1336R | CTCTGTTTTGGAAGAAGCTTTTGG | 86.32 |
| ASN102F | caaatttactggctgtttgatcc | 102 |
| ASN102R | ttatcgtcgacaagttaccaca | 102 |
| ASN103F | tggagctgttgccagagac | 110 |
| ASN103R | ctcagccaaggggtcaagta | 110 |
| dupssr31F | GATAGGAGTGCTGACGCTAA | 119 |
| dupssr31R | ATCCTGCTATAGAGTCCAGACTT | 119 |
| phi301654F | GAATGCATGCTTTTCAAGGAC | 132 |
| phi301654R | CGCACAGAGAGCAGAACG | 132 |
| bnlg1028F | AGGAAACGAACACAGCAGCT | 138.2 |
| bnlg1028R | TGCATAGACAAAACCGACGT | 138.2 |
| umc1113F | ATCATGCGTCATCACTCTCAGAAC | 147.9 |
| umc1113R | GCTGGAGCTAGCTGTAGTGTAGCA | 147.9 |

**Supplemental Table 4.**
**Recombination Mapping Data**

| Molecular marker | *3L* position (Mbp) | No. of chromatid types | | Estimated map units (cM) |
|---|---|---|---|---|
| | | A619 | A632 | |
| bnlg1505 | 147 | 191 | 5 | 0.025 |
| umc2265 | 159 | 195 | 1 | 0.005 |
| ANSSR161-2 | 161 | 196 | 0 | <0.005 |
| ANSSR163 | 163 | 196 | 0 | <0.005 |
| ANSSR165-2 | 165 | 191 | 5 | 0.025 |
| ANSSR166 | 166 | 190 | 6 | 0.030 |
| umc1596 | 167 | 188 | 8 | 0.041 |

**Supplemental Table 5.**
**List of Oligonucleotide Primers Used for Genomic DNA Amplifications and Sequencing**

| Primer Name | Pub. Primer Name | Region Targeted | Primer Locations | Sequence |
|---|---|---|---|---|
| DCL3c_OsF | DCL3_exon1-3_F | exon1,2,3 | 5'UTR | GCGTGGCAGCCGATTCCGCG |
| DCL3c_3R | DCL3_exon1-3_R | exon1,2,3 | exon3 | GCATTGCTGTGTGACGAGTT |
| DCL3-16F | DCL3_exon2-4_F | exon2,3,4 | intron1 | TGATCTCGCTTGACAAAGCA |
| DCL3-15R | DCL3_exon2-4_R | exon2,3,4 | intron4 | GAGCTAAACATCTGCTGTCAGAAA |
| DCL3-18F | DCL3_exon5_F | exon5 | exon4 | GAAGGCTCATAGCTGGCAAC |
| DCL3-18R | DCL3_exon5_R | exon5 | intron5 | CAAACGGGCAAAATAACAGAA |
| DCL3-21F | DCL3_exon6_F | exon6 | exon5 | GCCATCACGCAACTGGTAAC |
| DCL3-21R | DCL3_exon6_R | exon6 | intron6 | GGTGAAGATACAAAATCCATGGTAG |
| DCL3-24F | DCL3_exon7-8_F | exon7,8 | intron6 | TCCATAGAGGAATTCTGTGGGTA |
| DCL3-24R | DCL3_exon7-8_R | exon7,8 | intron8 | GACATCGATCTTAACAGCTTTGG |
| DCL3_8_F2 | DCL3_exon9_F | exon9 | exon8 | TGTTCCTTCTGCAAAAGAAGTG |
| DCL3_8_R1 | DCL3_exon9_R | exon9 | exon10 | TCAGCCAACCTTTTCTCTGG |
| DCL3_9_F2 | DCL3_exon10_F | exon10 | exon9 | TCACGGAAACGTCTCTCAAA |
| DCL3_9_R3 | DCL3_exon10_R | exon10 | exon11 | TGAGTTCAGAAGGGCATCAA |
| DCL3-33F | DCL3_exon11-13_F | exon11,12,13 | intron10 | TATGATGCATCGGTGCTGAC |
| DCL3-33R | DCL3_exon11-13_R | exon11,12,13 | intron13 | CCATCACAAGCATAAGTAGTTTAGAG |
| DCL3-37F | DCL3_exon14_F | exon14 | exon13 | TGATTTGCCGAGAACTACCC |
| DCL3-37R | DCL3_exon14_R | exon14 | exon15 | TTGAAACATGGCACTGGATG |
| DCL3-39F | DCL3_exon15_F | exon15 | intron14 | TGCAGGGTTCTAACCCATTT |
| DCL3-39R | DCL3_exon15_R | exon15 | exon16 | ATGCAGTTCTTTCCGTTTGG |
| DCL3-42F | DCL3_exon16_F | exon16 | intron15 | TGACACTGTCTTATTTCATTCTCAAA |
| DCL3-42R | DCL3_exon16_R | exon16 | exon17 | CCAATGAACAGTTTTCCAAACA |
| DCL3_16_F1 | DCL3_exon17_F | exon17 | intron16 | CCTGAAAGATTTCTTTTGGTTCTC |
| DCL3_16_R1 | DCL3_exon17_R | exon17 | intron17 | GCATATGAATATGTGAAAGATTTAGC |
| DCL3-48F | DCL3_exon18-19_F | exon18,19 | exon17 | TGACAAGAAAGAAGCAAAATTCC |
| DCL3-48R | DCL3_exon18-19_R | exon18,19 | exon19 | TGGCTGGCTAGCATTAGTGA |
| DCL3-1F | DCL3_exon19-20_F | exon19,20 | exon19 | TTGCTGATTCCCCTTGATTT |
| DCL3-1R | DCL3_exon19-20_R | exon19,20 | exon21 | GAGCAAGCCATCGACGAG |
| DCL3-4F | DCL3_exon21_F | exon21 | intron20 | AAAAGAGGCTGCATTTTGCT |
| DCL3-4R | DCL3_exon21_R | exon21 | exon22 | CAAGACTGCATCACCAAGGA |
| DCL3-6F | DCL3_exon22_F | exon22 | intron21 | GAACTTTGGCTAGTTTAAGCCTTTT |
| DCL3-6R | DCL3_exon22_R | exon22 | exon23 | TCCAAATCAAGTTTGCTGTCA |
| DCL3-8F | DCL3_exon23_F | exon23 | intron22 | TCCGACCCCAAAGATACTGT |
| DCL3-8R | DCL3_exon23_R | exon23 | exon24 | GCACGTTGTTGCAATGTTTT |
| DCL3-10F | DCL3_exon24-25_F | exon24,25 | intron23 | GTGAATCATGCTGCTGGTTG |
| DCL3-10R | DCL3_exon24-25_R | exon24,25 | exon26 | GCAGAATCCTGTGAGCTCTTC |
| DCL3-13F | DCL3_exon26-3'UTR_F | exon26,3'UTR | intron25 | GGTTGCAGTTGGCTTGTGTA |
| DCL3-13R | DCL3_exon26-3'UTR_R | exon26,3'UTR | 3' UTR | CATCAGACGGGAACGAGAGT |

### Supplemental Table 6.
### List of Oligonucleotide Primers Used for cDNA Amplifications and Sequencing

| Primer Name | Pub. Primer Name | Region Targeted | Primer Location | Sequence |
|---|---|---|---|---|
| DCL3c_OsF | DCL3_cDNA_5'UTR_F | 5'UTR-exon3 | 5'UTR | GCGTGGCAGCCGATTCCGCG |
| DCL3c_1F | DCL3_cDNA_exon1_F | exon2-exon3 | exon2 | AGAAGAGGGAATGCCAGGAT |
| DCL3c_Zm_F | DCL3_cDNA_exon2_F | exon2-exon3 | exon2 | GGAGACACGAATACTTTGAA |
| DCL3c_3R | DCL3_cDNA_exon3_R | 5'UTR-exon3 | exon3 | GCATTGCTGTGTGACGAGTT |
| DCL3-18F | DCL3_exon5_F | exon4-exon10 | exon4 | GAAGGCTCATAGCTGGCAAC |
| DCL3-21F | DCL3_exon6_F | exon5-exon10 | exon5 | GCCATCACGCAACTGGTAAC |
| DCL3_9_F2 | DCL3_exon10_F | exon9-exon15 | exon9 | TCACGGAAACGTCTCTCAAA |
| DCL3_8_R1 | DCL3_exon9_R | exon4-exon10 | exon10 | TCAGCCAACCTTTTCTCTGG |
| DCL3c_11_F | DCL3_cDNA_exon11_F | exon11-exon13 | exon11 | TGGGCTATATTTCACCAAAGC |
| DCL3-37F | DCL3_exon14_F | exon13-exon17 | exon13 | TGATTTGCCGAGAACTACCC |
| DCL3c_13_R | DCL3_cDNA_exon13_R | exon11-exon13 | exon13 | ACCCTCTTCTGCAACATCTG |
| DCL3-37R | DCL3_exon14_R | exon9-exon15 | exon15 | TTGAAACATGGCACTGGATG |
| DCL3CDNA-3F | DCL3_cDNA_exon16_F | exon16-exon18 | exon16 | TTCAGTTTCCCCTTGTGGAC |
| DCL3CDNA-3R | DCL3_cDNA_exon18_R | exon16-exon18 | exon18 | GGGTGGGTTTCAACACTAGC |
| DCL3-42R | DCL3_exon16_R | exon13-exon17 | exon17 | CCAATGAACAGTTTTCCAAACA |
| DCL3-48F | DCL3_exon18-19_F | exon17-exon19 | exon17 | TGACAAGAAAGAAGCAAAATTCC |
| DCL3-48R | DCL3_exon18-19_R | exon17-exon19 | exon19 | TGGCTGGCTAGCATTAGTGA |
| DCL3-1F | DCL3_exon19-20_F | exon19-exon24 | exon19 | TTGCTGATTCCCCTTGATTT |
| DCL3-1R | DCL3_exon19-20_R | exon19-exon21 | exon21 | GAGCAAGCCATCGACGAG |
| DCL3-4R | DCL3_exon21_R | exon19-exon22 | exon22 | CAAGACTGCATCACCAAGGA |
| DCL3c_24_F1 | DCL3_cDNA_exon24_F | exon24-3'UTR | exon24 | CACCACCTAGGCAAAAGGAA |
| DCL3-8R | DCL3_exon23_R | exon19-exon24 | exon24 | GCACGTTGTTGCAATGTTTT |
| DCL3-10R | DCL3_exon24-25_R | exon24-exon26 | exon26 | GCAGAATCCTGTGAGCTCTTC |
| DCL3-13R | DCL3_exon26-3'UTR_R | exon24-3'UTR | 3'UTR | CATCAGACGGGAACGAGAGT |

Yellow entries are identical to those used for genomic DNA amplifications (Supplemental Table 5).

### Supplemental Table 7.
### Locations of Domains Identified within ZmDCL3

| Sample | DEAD | Helicase-C | dsRNA_bind | PAZ | Rnase III a | Rnase III b |
|---|---|---|---|---|---|---|
| Zm Alignment_Pfam | 45-206 | 460-528 | 599-685 | 904-1036 | 1077-1211 | 1288-1399 |
| Zm_SMART | 38-242 | 444-531 | 680-699 | 868-1041 | 1059-1231 | 1267-1423 |
| Zm_PROSITE | 49-224 | 414-571 | 599-689 | 910-1017 | 1042-1211 | 1252-1400 |
| Zm_CCD | 56-206 | 409-528 | 599-678 | 884-1008 | 1061-1222 | 1268-1423 |
| Zm Envelope | 43-210 | 452-531 | 599-688 | 884-1039 | 1077-1211 | 1288-1400 |
| At Alignment_Pfam | 54-182 | 456-506 | | 838-979 | 1022-1157 | 1231-1339 |
| Os Alignment_Pfam | 39-104 | 392-462 | 533-614 | 835-973 | 1012-1143 | 1223-1334 |

| | | | | | | |
|---|---|---|---|---|---|---|
| At Envelope | 47-209 | 448-508 | | 824-982 | 1022-1157 | 1231-1340 |
| Os Envelope | 37-124 | 385-465 | 533-623 | 804-975 | 1012-1146 | 1223-1335 |

### Supplemental Table 8.
### List of *dcl* Genes Among the Grasses and Outgroups

| Organism | DCL | Locus |
|---|---|---|
| Arabidopsis thaliana | 1 | AT1G01040.2 |
| Arabidopsis thaliana | 2 | AT3G03300.1 |
| Arabidopsis thaliana | 3 | AT3G43920.2 |
| Arabidopsis thaliana | 4 | AT5G20320.1 |
| Aquilegia coerulea | 1 | Aquca_037_00286.1 |
| Aquilegia coerulea | 2 | Aquca_002_00892.1 |
| Aquilegia coerulea | 3a | Aquca_004_00102.1 |
| Aquilegia coerulea | 3b | Aquca_020_00511.1 |
| Aquilegia coerulea | 4 | Aquca_016_00137.1 |
| Oryza sativa | 1 | LOC_Os03g02970.1 |
| Oryza sativa | 2a | LOC_Os03g38740.1 |
| Oryza sativa | 2b | LOC_Os09g14610.1 |
| Oryza sativa | 3 | LOC_Os01g68120.1 |
| Oryza sativa | 5 | LOC_Os10g34430.1 |
| Oryza sativa | 4 | LOC_Os04g43050.1 |
| Brachypodium distachyon | 1 | Bradi1g77087.1 |
| Brachypodium distachyon | 2a | Bradi1g15440.1 |
| Brachypodium distachyon | 2b | Bradi1g21030.1 |
| Brachypodium distachyon | 3a | Bradi2g23187.2 |
| Brachypodium distachyon | 5 | Bradi3g29287.1 |
| Brachypodium distachyon | 3b | Bradi2g58270.1 |
| Brachypodium distachyon | 4 | Bradi5g15337.1 |
| Setaria italica | 1 | Si033853m |
| Setaria italica | 2 | Si033906m |
| Setaria italica | 3 | Si000027m |
| Setaria italica | 5 | Si039134m |
| Setaria italica | 4 | Si009168m |
| Sorghum bicolor | 1 | Chr1:72,107,638..72,118,572 |
| Sorghum bicolor | 2 | Sb01g015670.1 |
| Sorghum bicolor | 3 | Sb03g043355.1 |
| Sorghum bicolor | 5 | Sb01g018890.1 |
| Sorghum bicolor | 4 | Sb06g022180.1 |
| Zea mays | 1 | GRMZM2G040762_T01 |
| Zea mays | 2 | GRMZM2G301405_T01 |
| Zea mays | 3 | GRMZM5G814985_T01 |
| Zea mays | 5 | GRMZM2G413853_T01 |

| | | |
|---|---|---|
| Zea mays | 4 | GRMZM2G024466_T01 |
| Panicum virgatum | 1a | Pavirv00068715m |
| Panicum virgatum | 1b | Pavirv00059863m |
| Panicum virgatum | 2a | Pavirv00065721m |
| Panicum virgatum | 2c | Pavirv00017234m |
| Panicum virgatum | 3a | Pavirv00001779m |
| Panicum virgatum | 3b | Pavirv00022632m |
| Panicum virgatum | 3c | Pavirv00009554m |
| Musa acuminata | 1 | GSMUA_Achr8T12350_001 |
| Musa acuminata | 3a | GSMUA_Achr10T28110_001 |
| Musa acuminata | 3b | GSMUA_Achr10T28120_001 |
| Musa acuminata | 5 | GSMUA_Achr5T10880_001 |
| Musa acuminata | 4 | GSMUA_Achr9T22460_001 |
| Musa acuminata | 4 | GSMUA_Achr9T22470_001 |

Entries are color coded according to DCL clades (yellow = DCL1, Orange = DCL2, Green = DCL3 and DCL5, Blue = DCL4).

**Supplemental Table 9.**
**Summary Statistics of SBS Libraries**

| Progeny ID | Genotype | Total reads[1] | Genome matched[2] | Distinct genome matched[3] | Proportion distinct[4] |
|---|---|---|---|---|---|
| 080580 | *Rmr5 / rmr5-2* | 27,918,595 | 14,804,448 | 6,208,041 | 0.22 |
| 080580 | *rmr5-2 / rmr5-2* | 22,984,093 | 11,322,365 | 5,084,065 | 0.22 |

[1] Total reads >15 bp in length (after trimming of the 3′ adapter) sequenced.
[2] Total reads mapped to the 4a.53 AGPv1 maize genome, excluding r/t/sn/snoRNA.
[3] Total distinct reads, excluding r/t/sn/snoRNA, mapping to the 4a.53 AGPv1 found within the set.
[4] Number of distinct reads mapped to the B73 4a.53 AGPv1 divided by the total sequenced provides the proportion of distinct sequences.

**Supplemental Table 10.**
**Analysis of SBS Reads**

| Reads from SBS libraries | | | | | | | | |
|---|---|---|---|---|---|---|---|---|
| Size (nt) | Total genome mapping | | | | Unique genome mapping | | | |
| | *Rmr5 / rmr5-2* | | *rmr5-2 / rmr5-2* | | *Rmr5 / rmr5-2* | | *rmr5-2 / rmr5-2* | |
| 18 | 10381 | 0.07% | 32291 | 0.29% | 5650 | 0.09% | 26790 | 1% |
| 19 | 33000 | 0.22% | 298290 | 2.63% | 28026 | 0.45% | 204644 | 4% |
| 20 | 112187 | 0.76% | 907707 | 8.02% | 66868 | 1.08% | 497905 | 10% |
| 21 | 816631 | 5.52% | 3005237 | 26.54% | 246132 | 3.96% | 1143112 | 22% |
| 22 | 1257825 | 8.50% | 4878081 | 43.08% | 362134 | 5.83% | 1844187 | 36% |
| 23 | 1549700 | 10.47% | 922055 | 8.14% | 948254 | 15.27% | 512521 | 10% |
| 24 | 10800687 | 72.96% | 671920 | 5.93% | 4374056 | 70.46% | 402774 | 8% |
| 25 | 206407 | 1.39% | 322377 | 2.85% | 161588 | 2.60% | 218924 | 4% |
| 26 | 11349 | 0.08% | 136474 | 1.21% | 10281 | 0.17% | 107195 | 2% |
| 27 | 3135 | 0.02% | 76789 | 0.68% | 2642 | 0.04% | 63439 | 1% |
| 28 | 1807 | 0.01% | 45418 | 0.40% | 1327 | 0.02% | 39437 | 1% |
| 29 | 930 | 0.01% | 19585 | 0.17% | 740 | 0.01% | 17496 | 0% |
| 30 | 317 | 0.00% | 5108 | 0.05% | 264 | 0.00% | 4675 | 0% |
| 31 | 92 | 0.00% | 1033 | 0.01% | 79 | 0.00% | 966 | 0% |
| 32 | 0 | 0.00% | 0 | 0.00% | 0 | 0.00% | 0 | 0% |
| 33 | 0 | 0.00% | 0 | 0.00% | 0 | 0.00% | 0 | 0% |
| 34 | 0 | 0.00% | 0 | 0.00% | 0 | 0.00% | 0 | 0% |
| Totals: | 14804448 | 100.00% | 11322365 | 100.00% | 6208041 | 100.00% | 5084065 | 100% |
